# The histone chaperone FACT coordinates H2A.X-dependent signaling and repair of DNA damage

**DOI:** 10.1101/264952

**Authors:** Sandra Piquet, Florent Le Parc, Siau-Kun Bai, Odile Chevallier, Salomé Adam, Sophie E. Polo

## Abstract

Safeguarding cell function and identity following a genotoxic stress challenge entails a tight coordination of DNA damage signaling and repair with chromatin maintenance. How this coordination is achieved and with what impact on chromatin integrity remains elusive. Here, by investigating the mechanisms governing the distribution of H2A.X in mammalian chromatin, we demonstrate that this histone variant is deposited *de novo* at sites of DNA damage in a repair synthesis-coupled manner. Our mechanistic studies further identify the histone chaperone FACT (Facilitates Chromatin Transcription) as responsible for the deposition of newly synthesized H2A.X. Functionally, FACT potentiates H2A.X-dependent signaling of DNA damage and, together with ANP32E (Acidic Nuclear Phosphoprotein 32 Family Member E), orchestrates a H2A.Z/H2A.X exchange reaction that reshapes the chromatin landscape at repair sites. We propose that this mechanism promotes chromatin accessibility and helps tailoring DNA damage signaling to repair progression.

**HIGHLIGHTS:**

- H2A.X, but not H2A.Z, is deposited *de novo* at sites of DNA damage repair
- FACT promotes new H2A.X deposition coupled to repair synthesis
- FACT and ANP32E chaperones orchestrate H2A.Z/H2A.X exchange in damaged chromatin
- FACT stimulates H2A.X-dependent signaling of DNA damage

## INTRODUCTION

Cells are constantly exposed to genotoxic stress, and respond by activating dedicated DNA damage signaling and repair pathways that safeguard genome stability (Aguilera and García-Muse, 2013; Ciccia and Elledge, 2010; Hoeijmakers, 2009; Jackson and Bartek, 2009). DNA damage signaling consists in the activation of checkpoint kinase cascades upon DNA damage detection/processing to coordinate cell cycle progression with DNA repair (Lazzaro et al., 2009; Shaltiel et al., 2015). Adding another layer of complexity to these highly orchestrated cellular responses, damage signaling and repair machineries operate on chromatin substrates in eukaryotic cell nuclei, where DNA wraps around histone proteins (Luger et al., 2012). Importantly, chromatin landscapes, defined by specific patterns of histone variants (Buschbeck and Hake, 2017; Talbert and Henikoff, 2017), post-translational modifications (Bannister and Kouzarides, 2011) and various degrees of chromatin folding, convey epigenetic information that instructs cell function and identity through the regulation of gene expression programs (Allis and Jenuwein, 2016). While the importance of maintaining epigenome integrity is widely recognized, it is still poorly understood how this is achieved during the DNA damage response.

DNA damage signaling and repair indeed elicit profound chromatin rearrangements, challenging epigenome maintenance (Dabin et al., 2016), with a transient destabilization of chromatin organization, accompanied by DNA damage-induced changes in histone modifications (Dantuma and van Attikum, 2016). Chromatin structure is then restored concomitantly with the repair of DNA damage (Polo and Almouzni, 2015; Smerdon, 1991). It remains unclear whether chromatin restoration after damage is an entirely faithful process or if genotoxic stress responses alter the epigenetic landscape, leaving a signature of DNA damage repair. Although not fully characterized yet, restoration of chromatin at damage sites involves the deposition of newly synthesized histones, as shown at sites of UVA and UVC damage in human cells with the *de novo* deposition of H2A, H3.1 and H3.3 histone variants (Adam et al., 2013; Dinant et al., 2013; Luijsterburg et al., 2016; Polo et al., 2006). Mechanistically, new histone deposition employs dedicated histone chaperones (Hammond et al., 2017), including the H3 variant-specific chaperones HIRA (Histone Regulator A) and CAF-1 (Chromatin Assembly Factor-1), which are recruited to UV-damaged chromatin by early and late repair proteins, respectively (Adam et al., 2013; Polo et al., 2006). Less is known about the factors that govern H2A dynamics in damaged chromatin, apart from a role for the histone chaperone FACT in promoting H2A-H2B turnover at UVC damage sites (Dinant et al., 2013). Regarding the histone variant H2A.Z, its dynamic exchange at sites of DNA double-strand breaks in human cells involves the concerted action of the histone chaperone ANP32E and of the chromatin remodeling factors p400 and INO80 (Inositol-requiring 80)(Alatwi and Downs, 2015; Gursoy-Yuzugullu et al., 2015; Xu et al., 2012).

Here, we investigate how such repair-coupled chromatin rearrangements may leave an imprint on the epigenetic landscape and cross-talk with damage signaling by focusing on the H2A.X histone variant, which represents the ancestral form of H2A, conserved in all eukaryotes (Talbert and Henikoff, 2010). Making up only 10 to 25% of total H2A, H2A.X is nevertheless central to the DNA damage response, owing to a particular carboxyl-terminal Serine, in position 139 in mammals, that is targeted by DNA damage responsive kinases (Bonner et al., 2008; Rogakou et al., 1998). Importantly, H2A.X S139 phosphorylation spreads at a distance from the damage (Rogakou et al., 1999), which is key for amplifying the DNA damage signal through the coordinated recruitment of DNA damage checkpoint mediators (Altmeyer and Lukas, 2013; Smeenk and van Attikum, 2013). Best described in response to DNA double-strand breaks, this signaling cascade also operates following other types of genomic insults including UV irradiation. Indeed, UV damage processing triggers checkpoint signaling (Marini et al., 2006) by activating the ATR (ataxia telangiectasia-mutated and Rad3-related) kinase, which phosphorylates H2A.X (Hanasoge and Ljungman, 2007). This in turns recruits checkpoint mediators, including MDC1 (Mediator of DNA damage Checkpoint 1) (Marteijn et al., 2009).

Regardless of which type of DNA insult activates the H2A.X signaling cascade, a salient feature of H2A.X phosphorylation is that it takes place *in situ* in damaged chromatin (Rogakou et al., 1998). The original distribution of H2A.X in chromatin is thus a critical determinant of the damage response, as it will govern the distribution of the phosphorylated form, known as γH2A.X. A commonly held view is that H2A.X is phosphorylated at DNA damage sites but ubiquitously incorporated in chromatin, independently of DNA damage. However, recent ChIP-seq studies in mammalian cells have challenged this view by revealing a non-random distribution of H2A.X, with enrichments at active transcription start sites and sub-telomeric regions in activated human lymphocytes (Seo et al., 2014; 2012), and at extra-embryonic genes in mouse pluripotent stem cells (Wu et al., 2014). Yet, the mechanisms underpinning the non-random distribution of H2A.X in chromatin are unknown, as is their potential connection to the DNA damage response.

In this study, by investigating the mechanisms governing H2A.X distribution in chromatin during the repair of UVC damage in mammalian cells, we reveal that H2A.X histones are deposited *de novo* in damaged chromatin by the histone chaperone FACT concomitantly with repair synthesis. We also uncover a H2A.Z/H2A.X exchange reaction orchestrated by FACT and ANP32E chaperones, which reshapes the chromatin landscape by altering the histone variant pattern at UV damage sites. Functionally, these histone chaperones are key for mounting an efficient cellular response to DNA damage, with FACT potentiating H2A.X-dependent damage signaling.

## RESULTS

### *De novo* deposition of H2A histone variants at repair sites

In order to characterize H2A.X deposition pathways, we monitored *de novo* histone deposition using the SNAP-tag technology (Bodor et al., 2012) in human U2OS cell lines stably expressing SNAP-tagged H2A variants (Figure 1A and Figure S1 for a characterization of the cell lines). Specific labeling of newly synthesized histones combined with local UVC irradiation did not reveal any detectable accumulation of new H2A variants at UVC damage sites, contrary to what we had observed with newly synthesized H3.3 (Adam et al., 2013) (Figure 1B). We reasoned that this discrepancy might be due to differences in mobility between inner core histones, H3-H4, and outer core histones, H2A-H2B, the latter being more readily exchanged (Kimura and Cook, 2001; Louters and Chalkley, 1985), which may hinder the detection of local histone accumulations. Since outer core histone mobility is partly transcription-dependent (Jackson, 1990; Kimura and Cook, 2001), we tracked new histones in the presence of transcription inhibitors, Flavopiridol, DRB or α-amanitin (Bensaude, 2011) (Figures 1C and S2A). Note that short-term transcription inhibition reduces but does not abolish histone neosynthesis because of pre-existing mRNAs. Thus, we revealed new H2A.X accumulation at sites of UVC damage in the vast majority of cells (over 85%, Figure 1C-E), and we recapitulated our observations in mouse embryonic fibroblasts (Figure S3A-D). Importantly, new H2A.X accumulation at UVC damage sites was not an artifact of transcription inhibition as it was also detectable in the absence of transcription inhibitors upon exposure to higher UVC doses (Figure S2B). In contrast to new H2A.X, no significant enrichment was observed at damaged sites when staining for total H2A.X (Figure 1D), showing that this accumulation was specific for newly synthesized histones and most likely reflects histone exchange at damage sites. Noteworthy, *de novo* accumulation of H2A.X was also observed at sites of UVA laser micro-irradiation (Figure S2C), arguing that it is not unique to the UVC damage response.

**Figure 1.**
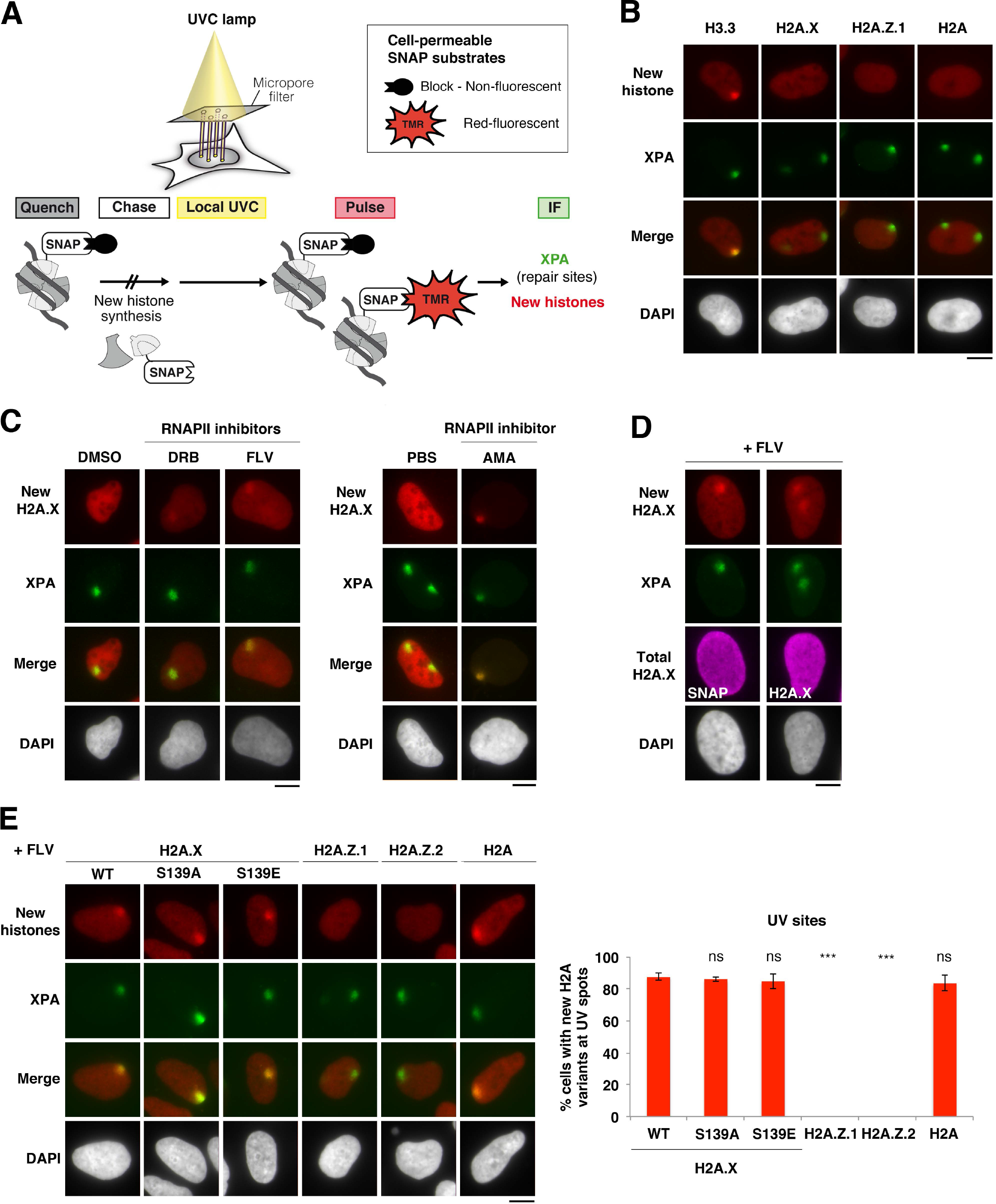
*De novo* accumulation of H2A variants at UVC damage sites. **(A)** Scheme of the assay for monitoring the accumulation of newly synthesized histones at UVC damage sites in cultured human cells stably expressing SNAP-tagged histones. Preexisting SNAP-tagged histones are quenched with a non-fluorescent substrate (block) so that only the histones neo-synthesized during the chase period are labeled with the red fluorescent substrate tetramethylrhodamine (TMR)-star during the pulse step. Local UVC damage is induced before the pulse step and damage sites are detected by immunostaining for the repair factor XPA (Xeroderma Pigmentosum, complementation group A). **(B)** New histone accumulation (red) at sites of UVC damage marked by the repair factor XPA (green) analyzed in U2OS cell lines stably expressing the indicated SNAP-tagged histone variants. **(C)** New H2A.X accumulation at sites of UVC damage marked by XPA analyzed 2 h after irradiation in U2OS cells stably expressing SNAP-tagged H2A.X and treated with the indicated RNAPII inhibitors (DRB; FLV: flavopiridol; AMA: α-amanitin; DMSO and PBS: vehicles). **(D)** New H2A.X accumulation at sites of UVC damage marked by XPA analyzed 2 h after irradiation in the presence of flavopiridol (+FLV) in U2OS cells stably expressing SNAP-tagged H2A.X. Total H2A.X are detected by immunostaining for SNAP or H2A.X. **(E)** *De novo* accumulation of H2A variants at sites of UVC damage marked by XPA analyzed in the presence of flavopiridol (+FLV) in U2OS cell stably expressing the indicated SNAP-tagged H2A variants. WT: wild-type, S139A: phospho-deficient mutant, S139E: phosphomimetic mutant. Percentages of cells accumulating new H2A variants at UV sites are shown on the graph. Error bars represent SD from two independent experiments. Scale bars, 10 µm. See also, Figures S1-S4.

In order to assess whether new H2A.X deposition at UVC damage sites was specific for the damage-responsive histone H2A.X, we extended our analyses to other H2A variants, namely canonical H2A and another replacement variant conserved in all eukaryotes, H2A.Z, considering both H2A.Z.1 and H2A.Z.2 forms, which display different dynamics in response to UVA laser damage in human cells (Nishibuchi et al., 2014). For this, we established U2OS cell lines that stably express comparable levels of SNAP-tagged H2A variants (Figure S1). Similar to what observed for H2A.X, we detected *de novo* accumulation of the canonical replicative variant H2A, but not of the replacement variants H2A.Z.1 and H2A.Z.2, at UVC damage sites (Figure 1E). Similar results were obtained without transcription inhibition (Figure S2D), pointing to a specific histone deposition mechanism that is not general to all H2A variants.

Collectively, these data reveal a transcription-independent deposition of newly synthesized H2A and H2A.X but not H2A.Z histone variants at repair sites.

### New H2A.X deposition at repair sites is independent of S139 phosphorylation

We next examined the importance of H2A.X S139 phosphorylation status for new H2A.X deposition at repair sites. We first verified that the SNAP-tag in C-terminal did not prevent H2A.X phosphorylation (data not shown) and we compared the dynamics of H2A.X wild-type to phospho-mimetic (S139E) and phospho-deficient (S139A) mutants. These experiments consistently showed that new H2A.X accumulation at sites of UVC damage repair occurred irrespective of H2A.X S139 phosphorylation status (Figure 1E). Since SNAP-tagged H2A.X proteins are expressed in the context of wild-type endogenous H2A.X, we used a complementary approach to address the importance of H2A.X phosphorylation: we treated cells with an ATR kinase inhibitor, which inhibited UV-induced H2A.X phosphorylation as reported (Hanasoge and Ljungman, 2007), but did not impair the *de novo* deposition of H2A.X at UVC-damage sites (Figure S4A). Thus, H2A.X phosphorylation in response to DNA damage is dispensable for the deposition of new H2A.X at sites of DNA damage repair.

### *De novo* deposition of H2A histone variants at replication foci

In parallel to our analysis of H2A.X dynamics at sites of DNA repair, we also monitored new H2A.X deposition in the absence of DNA damage. Indeed, we noticed that new H2A.X displayed a punctuate pattern in a subset of undamaged cells upon transcription inhibition. This deposition pattern corresponded to replication foci, as shown by co-staining with the replication-coupled histone chaperone CAF-1 (Figure 2A). We observed new H2A.X accumulation at replication foci throughout S phase in human U2OS cells treated with transcription inhibitors (Figure 2B). In mouse embryonic fibroblasts, transcription inhibitors could be omitted to visualize new H2A.X accumulation in replicating heterochromatin domains, which are poorly transcribed by nature (Figure S3E-F). Similar to H2A.X accumulation at repair sites, the enrichment observed at replication foci was specific for newly synthesized H2A.X (Figure 2C) and only detected for H2A and H2A.X, but not H2A.Z variants (Figure 2D).

**Figure 2.**
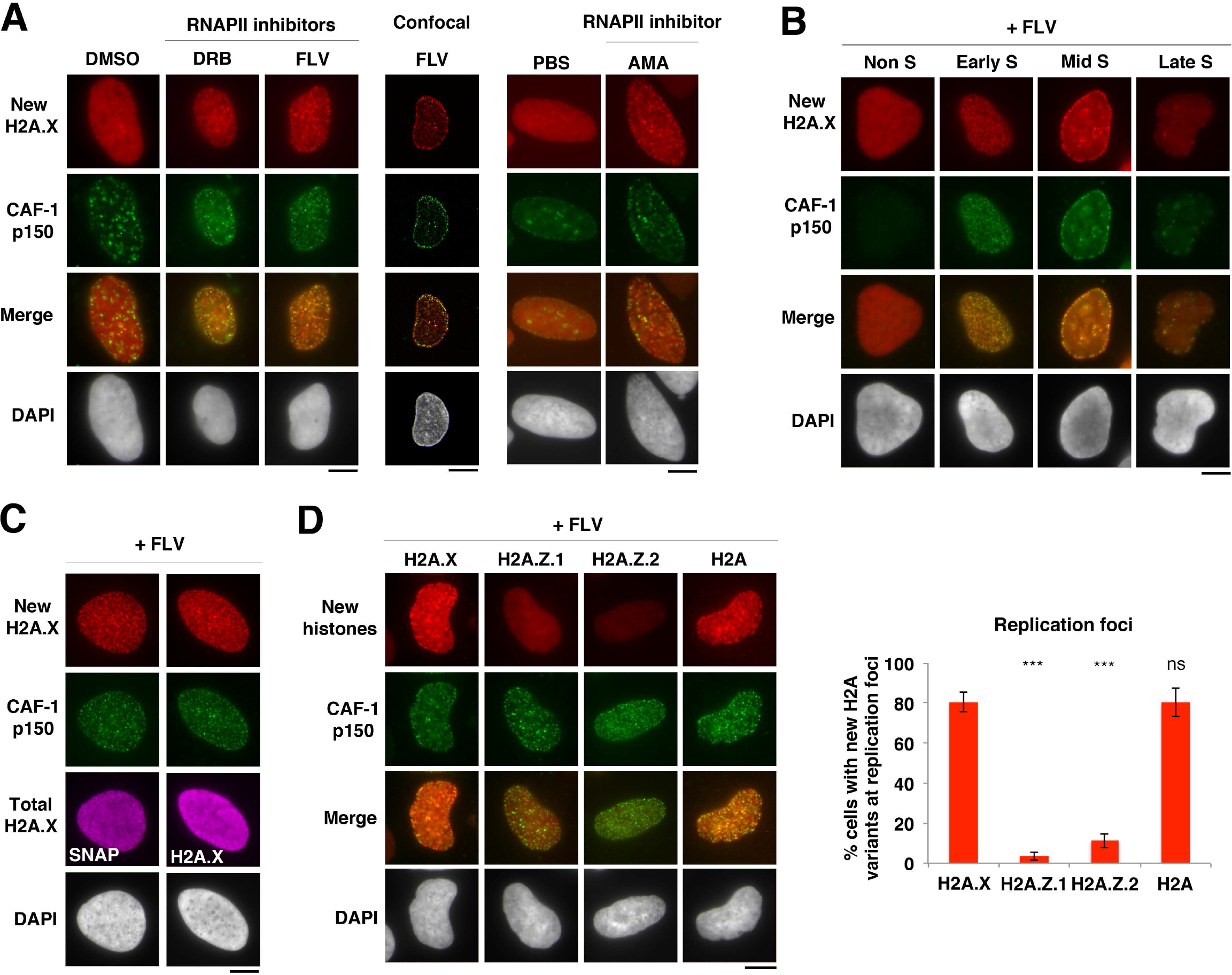
*De novo* accumulation of H2A variants at replication foci. **(A)** New H2A.X accumulation at sites of replication foci marked by CAF-1 p150 analyzed in U2OS cells stably expressing SNAP-tagged H2A.X and treated with the indicated RNAPII inhibitors (DRB; FLV: flavopriridol; AMA: α-amanitin; DMSO and PBS: vehicles). **(B)** New H2A.X accumulation at replication foci throughout S-phase analyzed in U2OS cells stably expressing SNAP-tagged H2A.X. Early, mid and late S phase cells are distinguished based on the pattern of replication foci. **(C)** New H2A.X accumulation at replication foci marked by CAF-1 p150 analyzed in U2OS cells stably expressing SNAP-tagged H2A.X. Total H2A.X is detected by immunostaining for SNAP or H2A.X. **(D)** *De novo* accumulation of H2A variants at replication foci marked by CAF-1 p150 analyzed in U2OS cell stably expressing the indicated SNAP-tagged H2A variants (H2A.X corresponds to the wild-type form). Percentages of cells accumulating new H2A variants at replication foci are shown on the graph. Error bars represent SD from two independent experiments. +FLV: flavopiridol. Scale bars, 10 µm. See also, Figures S1-S4.

Altogether, these findings demonstrate that newly synthesized H2A and H2A.X are deposited both at repair sites and at replication foci.

### New H2A.X deposition is coupled to replicative and repair synthesis

To gain insights into the mechanisms underlying new H2A.X deposition at repair sites and replication foci, we investigated a possible dependency on DNA synthesis, which is a common feature of replication and UV damage repair. We prevented repair synthesis by down-regulating the late repair factor XPG (Xeroderma Pigmentosum, group G), an endonuclease involved in the excision of the UVC-damaged oligonucleotide, which is a prerequisite for repair synthesis (Figure 3A). XPG knock-down did not impede damage detection by early repair factors including XPA but markedly reduced the accumulation of newly synthesized H2A.X at sites of UVC damage (Figure 3B) with no detectable effect on new H2A.X deposition at replication foci (data not shown). We confirmed that new H2A.X deposition at repair sites was dependent on the DNA synthesis machinery by knocking-down the DNA polymerase processivity factor PCNA (Proliferating Cell Nuclear Antigen, Figure 3C). In line with these observations at repair sites, we also uncovered a dependency on DNA synthesis for new H2A.X deposition occurring at replication foci. Indeed, when we inhibited replicative synthesis with aphidicolin, the focal patterns of new H2A.X deposition were strongly reduced in S phase cells, identified by EdU (Ethynyl-deoxyUridine) labeling (Figure 3D). These results indicate that the deposition of newly synthesized H2A.X at replication foci and UVC damage sites is dependent on replicative and repair synthesis machineries.

**Figure 3.**
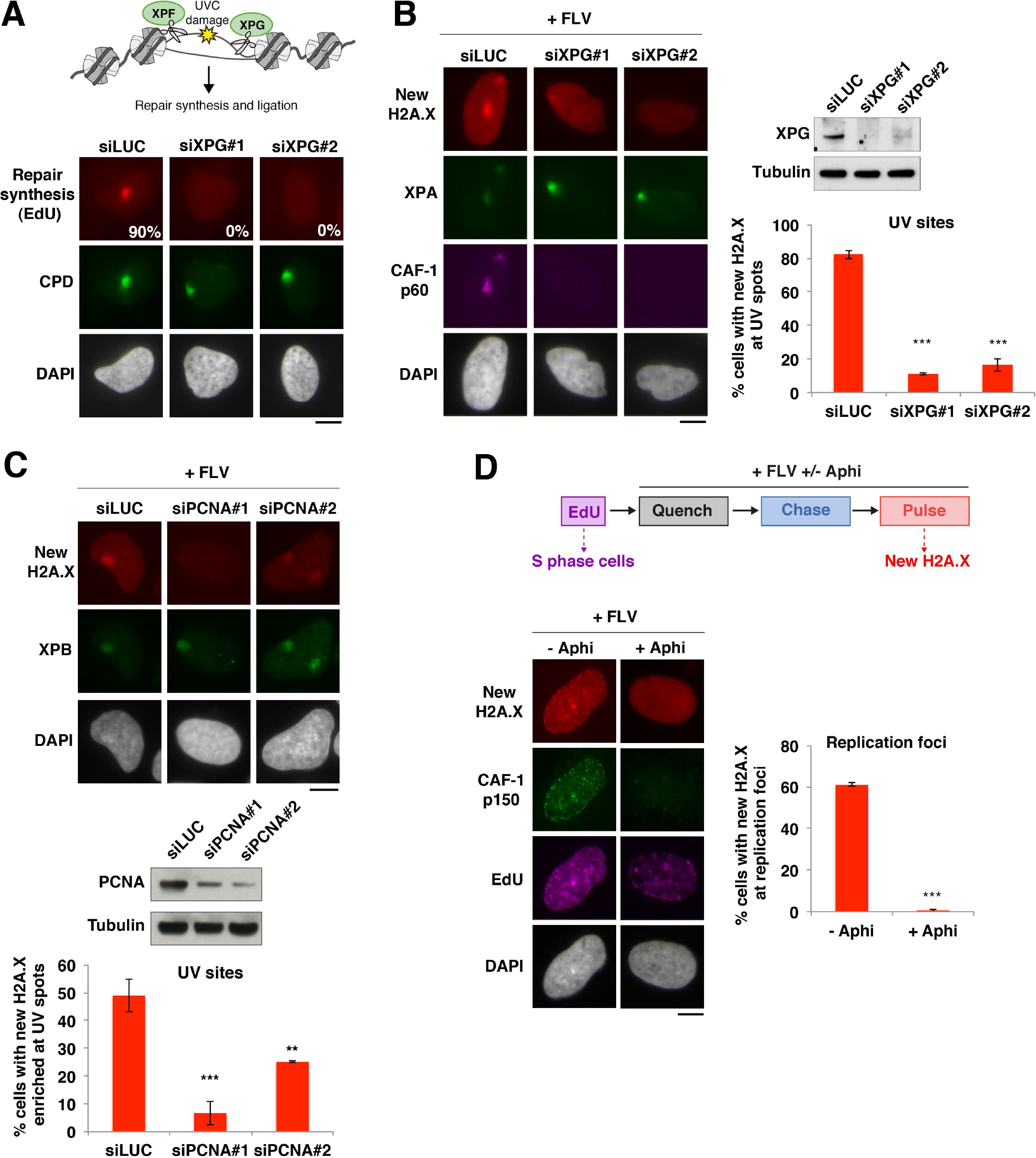
*De novo* accumulation of H2A.X at UV sites and replication foci is coupled to DNA synthesis. **(A)** Schematic representation of the role of the endonuclease XPG in late steps of UVC damage repair (top). Repair synthesis in U2OS cells treated with two different siRNAs targeting XPG (or siLUC: control) is measured by incorporation of 5-Ethynyl-2’-deoxyUridine (EdU) at UVC damage sites (Cyclobutane Pyrimidine Dimers, CPD). The percentage of cells showing repair synthesis is indicated (at least 200 cells were scored in 2 independent experiments). **(B)** New H2A.X accumulation at sites of UVC damage marked by the repair factor XPA analyzed 2 h after irradiation in the presence of flavopiridol (+FLV) in U2OS cells stably expressing SNAP-tagged H2A.X and transiently transfected with two different siRNAs targeting XPG (siLUC: control). Percentages of cells accumulating new H2A.X at UV sites are shown on the graph. The efficiency of XPG knock-down is verified by western-blot and by the lack of CAF-1 p60 recruitment to UV damage sites, which is coupled to repair synthesis. **(C)** New H2A.X accumulation at sites of UVC damage marked by the repair factor XPB (Xeroderma Pigmentosum, complementation group B) analyzed as in (B) in cells treated with two different siRNAs targeting PCNA (siLUC: control). Percentages of cells accumulating new H2A.X at UV sites are shown on the graph. The efficiency of PCNA knock-down is verified by western-blot. **(D)** Scheme of the experiment for testing the effect of replication inhibition by aphidicolin (Aphi) on new H2A.X accumulation at replication foci in U2OS H2A.X-SNAP cells treated with flavopiridol (+FLV). S-phase cells are labeled with EdU before Aphidicolin addition and new H2A.X accumulation is analyzed at replication foci marked by EdU. The efficiency of replication inhibition is shown by the lack of CAF-1 p150 recruitment to replication foci. Percentages of cells accumulating new H2A.X at replication foci are shown on the graph. Error bars on the graphs represent SD from at least two independent experiments. Scale bars, 10 µm. See also, Figure S4.

### New H2A.X deposition is not coupled to new H3 deposition

We next investigated whether new H2A.X deposition was coordinated with the deposition of newly synthesized H3 variants that occurs at replication and repair sites (Adam et al., 2013; Polo et al., 2006; Ray-Gallet et al., 2011). Knocking down the histone chaperone CAF-1, responsible for new H3.1 deposition at UV sites (Polo et al., 2006), did not significantly affect new H2A.X accumulation at sites of UVC damage repair (Figure S4B). Similar results were obtained upon down-regulation of the histone chaperone HIRA, which deposits new H3.3 at UV sites (Adam et al., 2013) (Figure S4B) and upon loss-of-function of both pathways simultaneously (data not shown). These data demonstrate that new H2A.X deposition occurs independently of new H3 deposition by CAF-1 and HIRA at UVC damage sites.

### The histone chaperone FACT promotes new H2A.X deposition at UV damage sites

To uncover the molecular determinants of new H2A.X accumulation at sites of UVC damage repair, we examined the effect of knocking down candidate H2A histone chaperones (Gurard-Levin et al., 2014; Hammond et al., 2017) and chromatin remodelers (Zhou et al., 2016). We focused on the histone chaperone FACT (Belotserkovskaya et al., 2003) and on the chromatin remodeler INO80 (Shen et al., 2000). Indeed, the latter is known to maintain the levels of chromatin-bound H2A.X in human cells (Seo et al., 2014) and to promote H2A deposition in yeast (Papamichos-Chronakis et al., 2006), while FACT incorporates H2A and H2A.X but not H2A.Z into chromatin (Heo et al., 2008) and stimulates H2A turnover at UVC damage sites in human cells (Dinant et al., 2013). Moreover, FACT associates with replisome components in human cells (Alabert et al., 2014; Tan et al., 2006), is involved in replication-coupled nucleosome assembly in yeast (Yang et al., 2016), and is critical for replisome progression through chromatin *in vitro* (Kurat et al., 2017). Interestingly, we observed that FACT accumulation at sites of DNA replication and UV damage repair exhibited dependencies to replicative synthesis and to late repair steps similar to new H2A.X deposition, FACT accumulation being impaired at repair sites by XPG and PCNA knock-downs and at replication foci upon aphidicolin treatment (Figures 4A-C and S5A). Furthermore, FACT trapping to chromatin by the intercalating agent curaxin CBL0137 (Gasparian et al., 2011; Safina et al., 2017) (Figure S5B) impaired FACT accumulation and new H2A.X deposition at UVC damage sites (Figure S5C-D). In line with these findings, siRNA-mediated down-regulation of both FACT subunits, but not of the remodeler INO80, led to a marked reduction of newly deposited H2A.X at UVC damage sites (Figure 4D). The decrease in new H2A.X deposition was also observed in the whole nucleus, most likely reflecting replication- and transcription-coupled deposition of H2A.X by FACT, and cannot be explained by reduced histone neo-synthesis as we verified that FACT knock-down does not inhibit nascent transcription (Figure S6A). *De novo* accumulation of H2A was similarly reduced in FACT-knocked-down cells (Figure S6B). These results establish that new H2A and H2A.X deposition at repair sites is promoted by the histone chaperone FACT.

**Figure 4.**
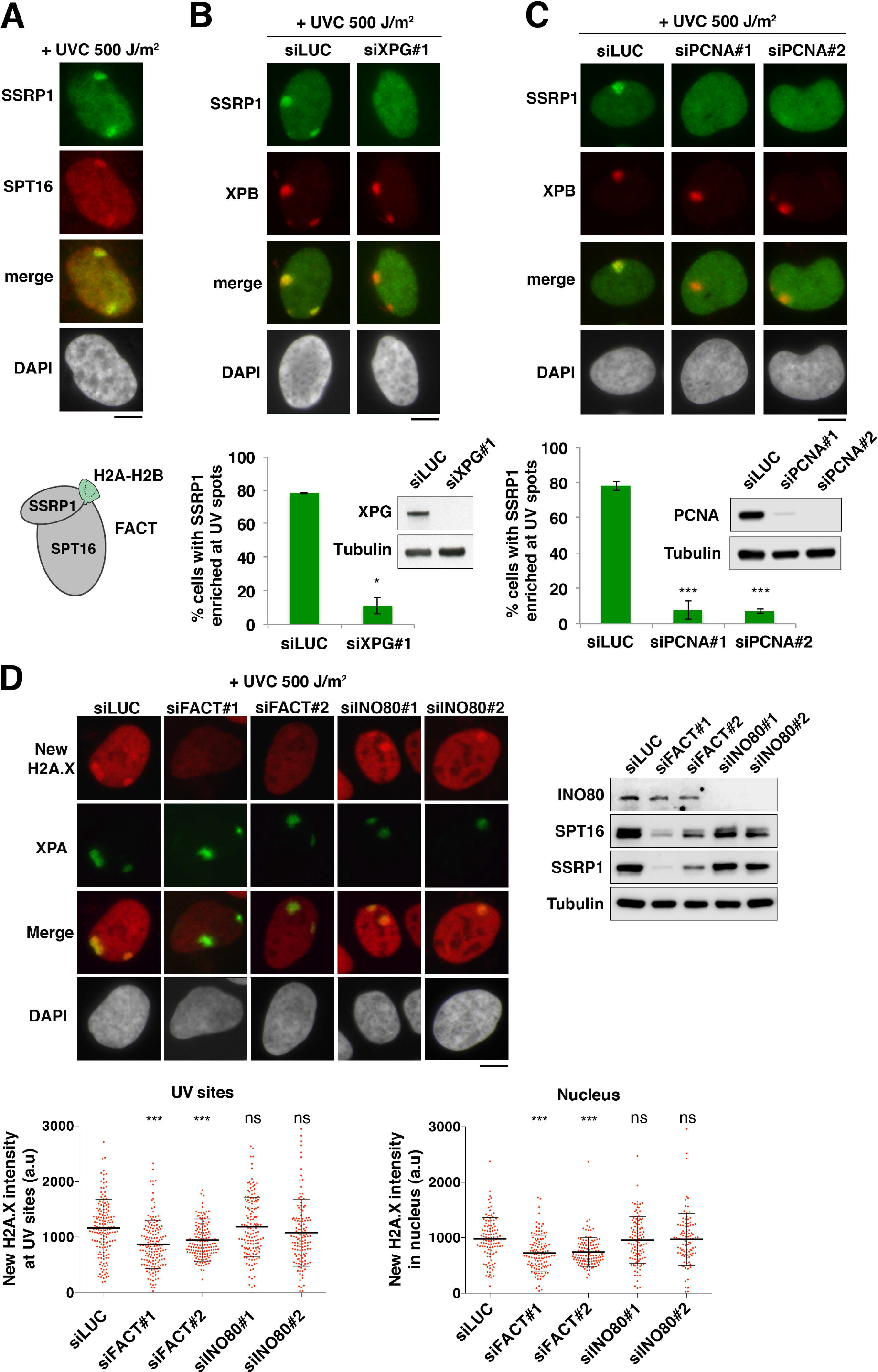
The histone chaperone FACT promotes new H2A.X deposition at UV damage sites. **(A)** Recruitment of FACT subunits, SSRP1 and SPT16, to repair sites 2 h after local UVC irradiation at 500 J/m^2^ in U2OS cells. The scheme represents FACT subunits bound to H2AH2B. **(B, C)** Recruitment of the histone chaperone FACT (SSRP1 subunit) to repair sites marked by XPB 2 h after local UVC irradiation at 500 J/m^2^ in U2OS cells treated with the indicated siRNAS (siLUC: control). Knock-down efficiency is controlled on the western-blot panels. Percentages of cells recruiting FACT to repair sites are shown on the graphs. Error bars represent SD from at least two independent experiments. **(D)** New H2A.X accumulation 1h30 after local UVC irradiation (500 J/m^2^) in U2OS H2A.X-SNAP cells treated with the indicated siRNAs (siLUC: control; siFACT: combination of siSPT16 and siSSRP1). Knock-down efficiencies are verified by western-blot. The intensity of new H2A.X signal (TMR fluorescence) at UV sites and in entire nuclei are shown on the graphs (bars: mean; error bars: SD from at least 100 cells). Similar results were obtained in three independent experiments and the results of a representative experiment are shown. Scale bars, 10 µm. See also, Figures S4-S6.

### H2A.Z/H2A.X exchange in UVC-damaged chromatin

The *de novo* deposition of H2A and H2A.X but not H2A.Z that we observed at sites of UV damage repair (Figure 1E) parallels the reported specificity of FACT towards the variants of H2A (Heo et al., 2008). We thus decided to examine whether this selective histone variant deposition by FACT may result in an altered histone variant pattern in UVC-damaged chromatin. Supporting this idea, when we stained for total histones, we observed that while H2A.X total levels were not detectably altered at sites of UVC irradiation (as also seen in Figure 1D), H2A.Z total levels were reduced by about 10% in damaged chromatin (Figure 5A-B). Importantly, the local reduction of H2A.Z was detectable with different antibodies targeting H2A.Z (amino- and carboxy-termini) and upon fluorescent labeling of total H2A.Z.1- and H2A.Z.2-SNAP (Figures 5A-B and S7A), ruling out the possibility of impaired detection due to post-translational modification of H2A.Z. Mechanistically, such selective depletion affecting the H2A.Z histone variant and not H2A.X at damage sites is unlikely to result from a local decompaction of chromatin. Furthermore, the reduction of H2A.Z in UVC-damaged chromatin is not a mere consequence of transcription inhibition at UV sites because transcription inhibition with flavopiridol does not significantly diminish the levels of chromatin-bound H2A.Z (Fig. S7B), suggesting that this histone variant is actively removed from UVC-damaged chromatin. When exploring the molecular determinants of H2A.Z removal, we found that the local depletion of H2A.Z occurred independently of FACT and INO80 but was dependent on the histone chaperone ANP32E (Figures 5B and S7C), previously characterized for its ability to remove H2A.Z from chromatin (Mao et al., 2014; Obri et al., 2014). Notably, ANP32E-mediated removal of H2A.Z was dispensable for FACT recruitment and new H2A.X deposition at UVC damage sites (Figure S7D-E). Thus, ANP32E and FACT chaperones independently orchestrate a histone variant exchange reaction in UVC-damaged chromatin resulting in the maintenance of H2A.X and the loss of H2A.Z.

**Figure 5.**
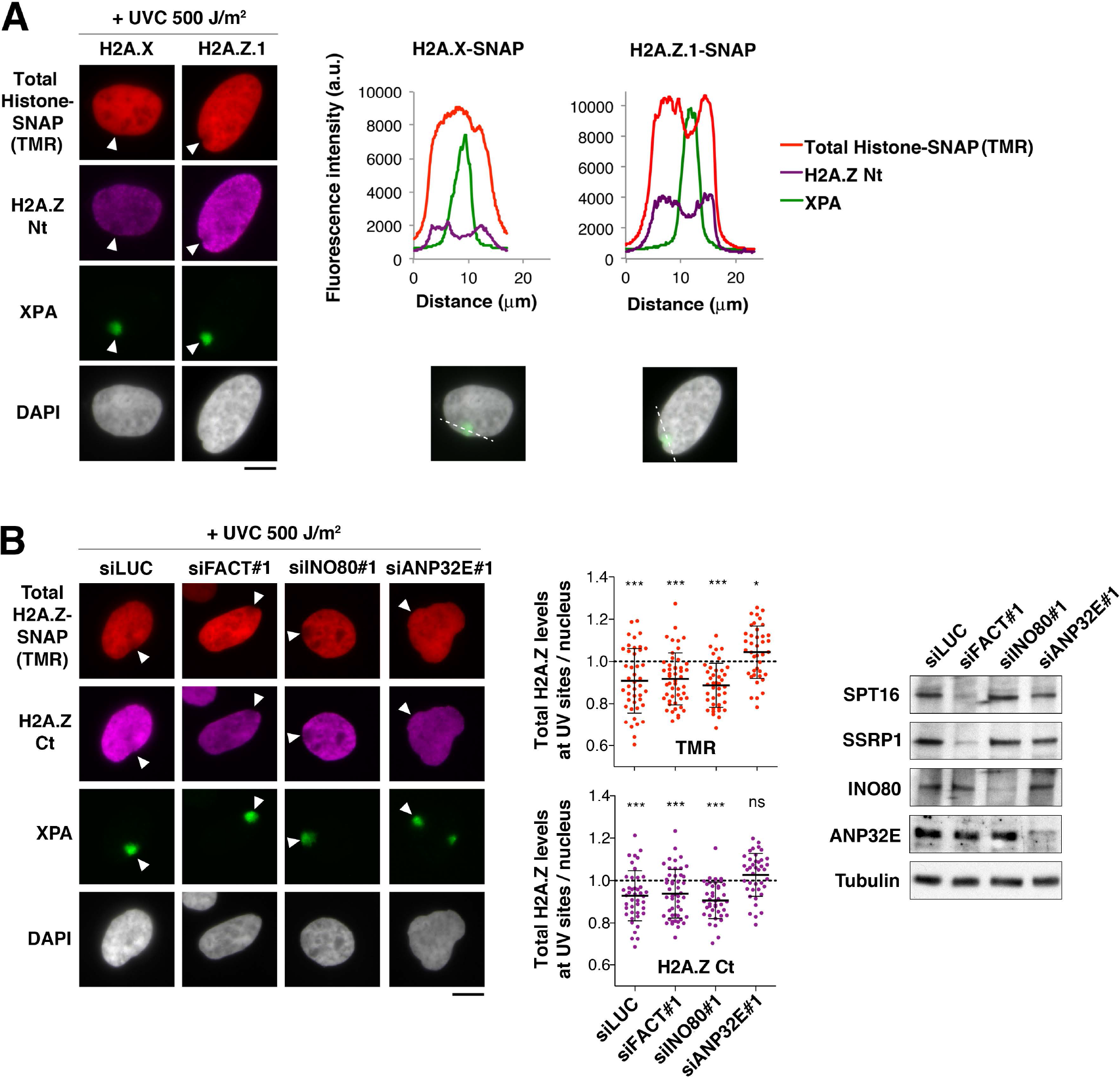
The histone chaperone ANP32E promotes H2A.Z removal from UV damage sites. **(A)** Distribution of total H2A.X and H2A.Z analyzed 1 h after local UVC irradiation (500 J/m^2^) in U2OS cells expressing the indicated SNAP-tagged histone variants. Total levels of SNAP-tagged histones are detected by a pulse with SNAP-Cell TMR-star (TMR) before cell fixation and total H2A.Z is also revealed by immunostaining with an H2A.Z-specific antibody recognizing H2A.Z amino-terminus (Nt). The arrowheads point to sites of UV irradiation. Fluorescence intensity profiles along the dotted lines are shown on the graphs. **(B)** Distribution of total H2A.Z 1 h after local UVC irradiation (500 J/m^2^) in U2OS H2A.Z.1-SNAP cells treated with the indicated siRNAs (siLUC, control; siFACT: combination of siSPT16 and siSSRP1). Total levels of SNAP-tagged H2A.Z are detected by a pulse with SNAP-Cell TMR-star (TMR) before cell fixation and total H2A.Z is revealed by immunostaining with an H2A.Z-specific antibody recognizing H2A.Z carboxy-terminus (Ct). Intensities in the red channel (TMR) were adjusted on a cell-by-cell basis because of variable levels of SNAP-tagged H2A.Z in the cell population. The arrowheads point to sites of UV irradiation. H2A.Z levels at UV damage sites relative to the whole nucleus are shown on the graphs (bars: mean; error bars: SD from at least 40 cells). The significance of H2A.Z loss or enrichment at UV sites is indicated (compared to a theoretical mean of 1, dotted line). Similar results were obtained in three independent experiments. Knock-down efficiencies are verified by western-blot. The specificity of H2A.Z Nt and Ct antibodies was verified by immunofluorescence upon H2A.Z knock-down by siRNA (data not shown). Scale bars, 10 µm. See also, Figures S7.

### FACT potentiates H2A.X-dependent signaling of DNA damage

Having identified FACT and ANP32E as key factors for H2A.Z/H2A.X exchange in damaged chromatin, we set out to determine the functional relevance of this mechanism. First, we observed that both FACT and ANP32E knock-downs confer increased sensitivity of cells to UVC damage (Figure 6A), suggesting that the histone variant exchange orchestrated by these two chaperones may be functionally important for the cells to mount an efficient DNA damage response. We further tested whether down-regulation of FACT or ANP32E might affect UVC damage repair. We observed that FACT and ANP32E knock-downs did not impair the recruitment of the repair factor XPA (Xeroderma Pigmentosum, group A) to UVC damage sites (Figures 4D, 5A-B, S6B, and S7A,C,E). In addition, we found no significant effect of FACT loss-of-function on repair synthesis (Figure S8A), in line with previous data (Dinant et al., 2013), and on the timely removal of UV photoproducts (Cyclobutane Pyrimidine Dimers, Figure S8B), ruling out any major effect of this histone chaperone on UVC damage repair. Similarly, preventing new H2A.X synthesis by siRNA (Figure S8C) did not detectably impair UVC damage repair synthesis (Figure S8D), arguing that FACT-mediated deposition of H2A.X in damaged chromatin does not impact DNA damage repair. Likewise, ANP32E was dispensable for repair synthesis at sites of UVC damage (Figure S8E). Together, these data do not reveal any significant contribution of FACT and ANP32E chaperones to the repair of UVC damage.

**Figure 6.**
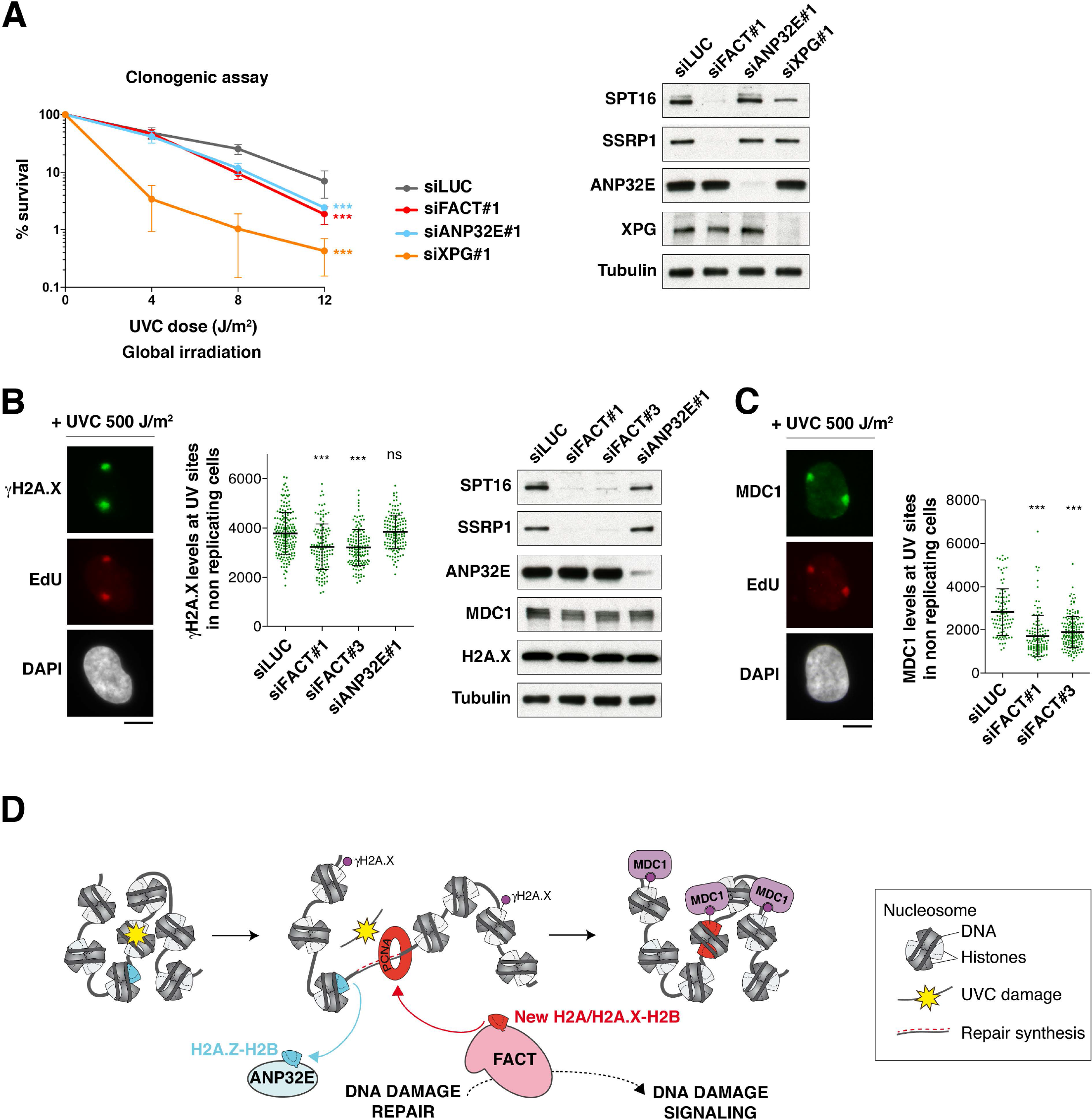
FACT potentiates H2A.X-dependent signaling of DNA damage. **(A)** Clonogenic survival of U2OS cells treated with the indicated siRNAs (siLUC, control) in response to global UVC irradiation. Error bars on the graph represent SD from four independent experiments. Knock-down efficiencies are verified by western-blot. **(B, C)** Quantification of γH2A.X and MDC1 levels (green) at sites of UVC damage repair (EdU spots, red) 2 h after local UVC irradiation (500 J/m^2^) in U2OS cells treated with the indicated siRNAs (siLUC, control). EdU was added to the culture medium immediately after UVC irradiation and left until cell fixation. Replicating cells showing pan-nuclear EdU staining were excluded from the analysis. Results from a representative experiment are shown (bars: mean; error bars: SD from at least 99 UV spots). Similar results were obtained in three independent experiments. Knock-down efficiencies are verified by western-blot. Scale bars, 10 µm. **(D)** Model for chromatin reshaping during UVC damage repair through the concerted activities of ANP32E and FACT histone chaperones promoting H2A.Z/H2A.X histone variant exchange. ANP32E removes H2A.Z from damaged chromatin (blue), which may enhance chromatin accessibility. FACT promotes new H2A.X deposition coupled to repair synthesis (red) and potentiates DNA damage signaling (purple), thus contributing to coordinate signaling and repair of DNA damage. The resulting distribution of H2A.X in chromatin reflects DNA damage experience. See also, Figure S8.

We next investigated a potential impact of these chaperones on DNA damage signaling after UVC irradiation. In particular, we hypothesized that FACT-mediated deposition of new H2A.X could potentiate damage signaling by bringing in more phosphorylation targets for DNA damage-responsive kinases or contribute to turn off damage signaling at the end of repair by replacing γH2A.X by unphosphorylated H2A.X and H2A. To avoid misinterpretation of the data due to the interference of FACT knock-down with DNA replication, we restricted our analysis to non-replicating cells based on staining with EdU (Ethynyl-deoxyUridine). Thus, we noticed that γH2A.X levels were reduced by 30 to 35% at sites of UVC damage repair in FACT-depleted cells (Figure 6B), while H2A.X total levels did not show any measurable decrease as assessed in total cell extracts and also at damage sites (Figure 6B and data not shown). Moreover, this reduction in γH2A.X levels did not result from lower damage infliction or from a diminished repair response because repair synthesis and repair factor recruitment were not significantly affected in the same cells (data not shown). In contrast to FACT loss-of-function, ANP32E depletion had no inhibitory effect on γH2A.X levels at UVC damage sites (Figure 6B). We also analyzed the recruitment to UVC-damaged chromatin of downstream damage signaling factors including the checkpoint mediator MDC1. In line with the reduced levels of γH2A.X, MDC1 accumulation at UVC damage sites also showed 30 to 40% reduction in FACT knocked-down cells while MDC1 total levels remained unaffected (Figure 6B-C). Collectively, these results establish that FACT and ANP32E histone chaperones are critical for the cells to mount an efficient DNA damage response with FACT stimulating H2A.X-dependent signaling of DNA damage.

## DISCUSSION

By analyzing the dynamics of H2A histone variants in damaged chromatin, we have uncovered a H2A.Z/H2A.X exchange reaction controlled by two histone chaperones, ANP32E and FACT (Figure 6D). Chromatin restoration after UVC damage repair thus entails reshaping of the chromatin landscape through a change in histone variant pattern. We also reveal that H2A.X is deposited *de novo* at sites of DNA damage repair, a process which should be integrated in the DNA damage response, in addition to the well-characterized phosphorylation of H2A.X. We propose that repair-coupled deposition of new H2A.X constitutes an important additional regulation level for fine-tuning the DNA damage response. By ensuring that the phosphorylation target for DNA damage checkpoint kinases is present in sufficient amount and at the right place, new H2A.X deposition may allow an efficient and timely response to genotoxic insults in chromatin regions that are susceptible to DNA damage.

### New H2A.X deposition at sites of DNA synthesis during replication and repair

In our study, we uncover a selective deposition of newly synthesized H2A variants at sites of UVC damage repair, with H2A and H2A.X being deposited *de novo* in damaged chromatin while H2A.Z is not. This parallels our previous findings on H3 variants showing that H3.1 and H3.3 are deposited *de novo* in UVC-damaged chromatin in contrast to CENP-A (Adam et al., 2013; Polo et al., 2006). Nevertheless, we demonstrate that new H2A.X deposition is not coupled with new H3.1/H3.3 deposition at repair sites. These results are in line with pioneering observations that newly synthesized H2A and H3 do not deposit within the same nucleosomes in replicating cells (Jackson, 1987), and suggest that new H2A.X-H2B associate with parental H3-H4 in repaired chromatin. We have also observed a selective deposition of H2A.X and H2A but not H2A.Z variants on replicating DNA, which is consistent with a recent study in mouse pericentric heterochromatin domains showing a replication-dependent deposition of new H2A and not H2A.Z (Boyarchuk et al., 2014), and with recent proteomic analyses of histone variants associated with newly replicating DNA in human cells (Alabert et al., 2015). The coupling between new H2A.X deposition and DNA replication is intriguing considering that H2A.X is not a prototypical replicative histone variant. Indeed, H2A.X is synthesized at basal levels throughout the cell cycle in mammalian cells, in a replication-independent manner (Wu and Bonner, 1981). Nevertheless, the lack of introns and the existence of a non-polyadenylated form of H2A.X mRNA - both characterizing replicative histone transcripts - place H2A.X in a unique position between replicative and replacement histone variants (Mannironi et al., 1989). Functionally, the deposition of newly synthesized H2A.X at replication foci may ensure that H2A.X is not diluted out during replication, which would happen if only the *bona fide* replicative variant H2A was deposited. Such replication-coupled deposition of H2A.X could also explain the increase in soluble H2A.X detected in human cells treated with replication inhibitors (Liu et al., 2008). Furthermore, the *de novo* deposition of H2A.X that we have uncovered at repair sites may provide a molecular basis for the H2A.X hotspots found in chromatin regions that are susceptible to endogenous damage (Seo et al., 2014). We can speculate that the observed enrichment of H2A.X may be a consequence of recurrent genomic insults. H2A.X could thus mark chromatin regions that are more susceptible to damage to facilitate subsequent damage responses. Noteworthy, the H2A.X protein was also shown to be rapidly stabilized following DNA damage, contributing to a local enrichment at repair sites (Atsumi et al., 2015). However, this stabilization required H2A.X phosphorylation on Ser139, while we found that new H2A.X deposition at repair sites occured independently of H2A.X phosphorylation. H2A.X protein stabilization and *de novo* deposition thus appear to be two distinct and complementary mechanisms contributing to the distribution of H2A.X in chromatin, which reflects DNA damage experience.

### Role of the histone chaperone FACT in new H2A.X deposition

We have identified FACT as the responsible histone chaperone for new H2A and H2A.X deposition at repair sites. Interestingly, FACT also promotes the deposition of another H2A variant, macroH2A1.2, at sites of replication stress in mammalian cells (Kim et al., 2017). However, FACT does not incorporate H2A.Z in chromatin (Heo et al., 2008), which is consistent with the lack of new H2A.Z deposition at sites of DNA synthesis. We demonstrate that FACT-mediated deposition of new H2A.X at damage sites is concomitant with repair synthesis and dependent on the DNA polymerase sliding clamp PCNA. This is reminiscent of the recruitment of CAF-1 to damaged DNA (Moggs et al., 2000), putting forward PCNA as a platform for histone chaperone recruitment to damaged chromatin. It will be interesting to examine whether FACT directly associates with PCNA. This is an attractive hypothesis considering that the FACT subunit SSRP1 (Structure-Specific Recognition Protein 1) harbors a non-canonical PIP (PCNA interacting protein) box (Mailand et al., 2013). Alternatively, FACT interaction with PCNA could be mediated by PCNA-binding factors like XRCC1 (X-ray repair cross-complementing protein 1)(Fan et al., 2004), which contributes to UVC damage repair (Moser et al., 2007), and was recently reported to associate with the FACT complex subunit SSRP1 (Gao et al., 2017). In addition, other factors associated with the DNA synthesis machinery and shown to cooperate with FACT, like MCM (MiniChromosome Maintenance) proteins (Tan et al., 2006), could also contribute to FACT recruitment to sites of UV damage repair.

### Impact of new H2A.X deposition on DNA damage signaling and repair

Although H2A.X is deposited *de novo* at sites of UVC damage repair, we have observed that this histone variant is not required for the repair of UVC lesions. In this respect, our findings are in line with previous studies in mouse embryonic stem cells knocked-out for H2A.X, which display increased genomic instability but are not particularly sensitive to UV irradiation (Bassing et al., 2002). Similar to H2A.X, the histone chaperone FACT is dispensable for repair synthesis at UV damage sites. This contrasts with FACT requirement for replicative DNA synthesis (Abe et al., 2011; Kurat et al., 2017; Okuhara et al., 1999), suggesting that FACT-mediated chromatin disassembly/re-assembly activity is more critical for replication fork progression through chromatin than for DNA synthesis *per se*.

Besides an effect on DNA repair, we also tested whether FACT-mediated deposition of new H2A.X in damaged chromatin may help coordinate DNA repair synthesis with damage signaling. Interestingly, the histone chaperones ASF1 and CAF-1 are required for checkpoint termination in response to DNA double-strand breaks (Chen et al., 2008; Diao et al., 2017; Kim and Haber, 2009), suggesting that the re-establishment of nucleosomal arrays contributes to turning off DNA damage signaling after repair of DNA damage. FACT in contrast enhances γH2A.X levels at sites of UV damage repair, thus potentiating DNA damage signaling. Although we cannot rule our that other FACT-dependent activities may be involved, it is tempting to speculate that FACT stimulates damage signaling through new H2A.X deposition. While this does not lead to any measurable increase in H2A.X total levels at repair sites, it may be enough to help amplify the γH2A.X signal. The repair-coupled deposition of H2A.X may also be functionally important for efficient signaling during a subsequent damage response by maintaining critical levels of H2A.X in chromatin regions susceptible to DNA damage.

### H2A.Z/H2A.X exchange at sites of DNA damage repair

Not only have we shown that H2A.X is deposited *de novo* at sites of UV damage repair but we have also observed that this is paralleled by the removal of H2A.Z, resulting in a local change in histone variant pattern. It still remains to be determined if the histone variant exchange takes place at the level of the nucleosome, H2A.X replacing H2A.Z, or at the level of the damaged chromatin domain. H2A.Z is removed from UVC-damaged chromatin by the histone chaperone ANP32E, showing interesting similarities with what has been reported in response to UVA laser damage (Gursoy-Yuzugullu et al., 2015). However, in this case ANP32E activity results in H2A.Z returning to basal levels in damaged chromatin after a transient accumulation while we observe a local loss of H2A.Z at UVC damage sites. We did not find any significant role for the remodeler INO80 in H2A.Z removal from UVC-damaged chromatin, in contrast to what observed at sites of UVA laser damage in human cells (Alatwi and Downs, 2015), suggesting that different types of DNA lesions may engage different chromatin remodeling machineries.

We have shown that ANP32E increases cell survival to UVC irradiation, suggesting that ANP32E-mediated removal of H2A.Z from UVC-damaged chromatin may be of functional importance. Given that H2A.Z promotes chromatin folding *in vitro* (Fan et al., 2002; 2004), the local depletion of H2A.Z could contribute to the chromatin relaxation observed at UVC damage sites (Adam et al., 2016; Luijsterburg et al., 2012). In line with this hypothesis, H2A.Z removal by ANP32E is required for histone H4 acetylation at sites of DNA double-strand breaks (Gursoy-Yuzugullu et al., 2015). Future studies will help decipher the functional contribution to the UV damage response of H2A.Z displacement by ANP32E and dissect potential crosstalks with other pathways controlling histone variant dynamics in damaged chromatin.

### Dynamics of H2A variants at repair sites and transcription regulation

While we have shown that the *de novo* deposition of H2A and H2A.X at repair sites was not transcription-mediated, it could impact transcription regulation in damaged chromatin. In particular, given the reported function of FACT in regulating transcription recovery after UVC damage in human cells (Dinant et al., 2013), FACT-mediated deposition of new H2A variants in UVC-damaged chromatin could facilitate the coordination of repair synthesis with transcription restart. Tipping the balance towards H2A and H2A.X as opposed to H2A.Z could also help keeping transcription in check and avoiding transcription interference with repair because H2A.Z was proposed to poise genes for transcription activation (Zhang et al., 2005) and to promote unscheduled/cryptic transcription when mislocalized in yeast cells (Jeronimo et al., 2015). It is thus tempting to speculate that H2A.Z/H2A.X histone variant exchange at UV sites may contribute to mitigate transcription-repair conflicts by maintaining transcription inhibition during repair synthesis. In this respect, it is worth mentioning that FACT was shown to limit the formation of RNA:DNA hybrids in yeast and human cells (Herrera-Moyano et al., 2014). It would thus be of major interest to investigate whether this function of FACT relies on the *de novo* deposition of H2A histone variants on newly synthesized DNA.

New H2A.X deposition in damaged chromatin could also have a broader impact on transcription regulation. Indeed, several recent studies have uncovered a critical role for H2A.X in regulating gene expression patterns during cell differentiation. In particular, H2A.X is involved in silencing of extra-embryonic genes in mouse embryonic stem cells (Wu et al., 2014), and also represses genes governing the epithelial-mesenchymal transition in human cells (Weyemi et al., 2016). Therefore, alterations in H2A.X distribution as a consequence of DNA damage experience may impact cell fate determination via the rewiring of transcriptional programs, thus contributing to the reported effect of DNA damage repair on cell differentiation and reprogramming (Rocha et al., 2013).

## METHOD DETAILS

### Cell culture and drug treatments

All U2OS (American Type Culture Collection ATCC HTB-96, human osteosarcoma, female) and NIH/3T3 (ATCC CRL-1658, mouse embryonic fibroblast, male) cell lines were grown at 37°C and 5% CO_2_ in Dulbecco’s modified Eagle’s medium (DMEM, Invitrogen) supplemented with 10% fetal calf serum (EUROBIO), 100 U/mL penicillin and 100 µg/mL streptomycin (Invitrogen) and the appropriate selection antibiotics (Table 1). Transcription inhibition was performed either by adding 5,6-Dichlorobenzimidazole 1-beta-Dribofuranoside (DRB, 100 µM final concentration, Sigma-Aldrich), alpha-Amanitin (AMA, 20 µg/mL final concentration, Sigma-Aldrich) or Flavopiridol hydrochloride hydrate (FLV, 10 µM final concentration, Sigma-Aldrich) to the culture medium at 37°C 4 h before harvesting the cells. Inhibition of replicative synthesis was performed by adding Aphidicolin (Aphi, 1 µg/mL final concentration, Sigma-Aldrich) to the culture medium at 37°C 5 h before harvesting the cells. For ATR inhibition, cells were incubated with 2 µM ATR inhibitor AZ20 (Selleckchem) 1h before subsequent cell treatment. For FACT trapping, cells were incubated with 2 µM CBL0137 (Bertin Pharma) 15 min before subsequent cell treatment.

**Table 1.** Stable cell lines. Antibiotics: G418 (100 µg/mL for U2OS, 500 µg/mL for NIH/3T3, Euromedex), Hygromycin (200 µg/mL, Euromedex).

| Cell line (reference if not generated in this study) | Selection antibiotics |
| --- | --- |
| U2OS H2A.X-SNAP WT form | G418 |
| U2OS H2A.X-SNAP S139A mutant | G418 |
| U2OS H2A.X-SNAP S139E mutant | G418 |
| U2OS H2A.Z.1-SNAP | G418 |
| U2OS H2A.Z.2-SNAP | G418 |
| U2OS H2A-SNAP | G418 |
| U2OS H3.3-SNAP (Dunleavy et al., 2011) | G418 |
| NIH/3T3 H2A.X-SNAP + GFP-DDB2 | G418 + Hygromycin |

### SNAP-tag labeling of histone proteins

For specific labeling of newly-synthesized histones, pre-existing SNAP-tagged histones were first quenched by incubating cells with 10 µM non-fluorescent SNAP reagent (SNAP-Cell Block, New England Biolabs) for 30 min (quench) followed by a 30 min-wash in fresh medium and a 2 h-chase. The SNAP-tagged histones neo-synthesized during the chase were then pulse-labeled by incubation with 2 µM red-fluorescent SNAP reagent (SNAP-Cell TMR-Star, New England Biolabs) for 15 min followed by 1h to 1h30-wash in fresh medium. Cells were pre-extracted with Triton detergent before fixation (see Immunofluorescence section for details). If cells were subject to local UVC irradiation, irradiation was performed immediately before the pulse. When transcription inhibitors were used, they were added to the medium at the quench step and kept throughout the experiment. For total labeling of SNAP-tagged histones, the quench step was omitted and cells were pulsed with SNAP-Cell TMR-Star immediately before harvesting.

### siRNA and plasmid transfections

siRNAs purchased from Eurofins MWG Operon (Table 2) were transfected into cells using Lipofectamine RNAiMAX (Invitrogen) following manufacturer’s instructions. The final concentration of siRNA in the culture medium was 50 nM. Cells were analyzed and/or harvested 48 to 72 h post-transfection. For stable cell line establishment, cells were transfected with plasmid DNA (1 µg/ml final) using Lipofectamine 2000 (Invitrogen) according to manufacturer’s instructions 48 h before antibiotic selection of clones. All constructs were verified by direct sequencing and/or restriction digests. Cloning details and primer sequences (Sigma-Aldrich) are available upon request. Plasmids are described in Table 3.

**Table 2.**
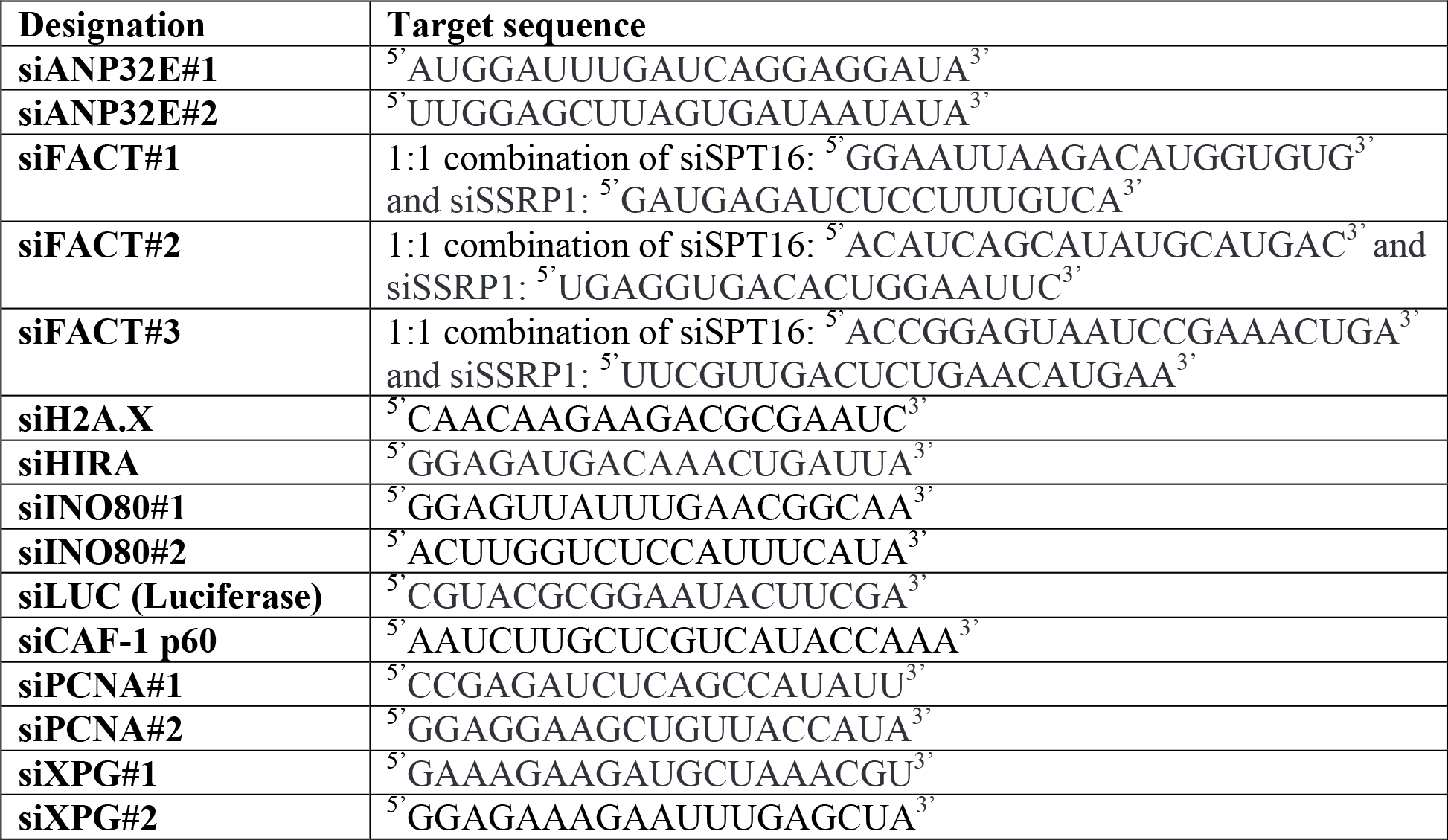
siRNA sequences.

**Table 3.**
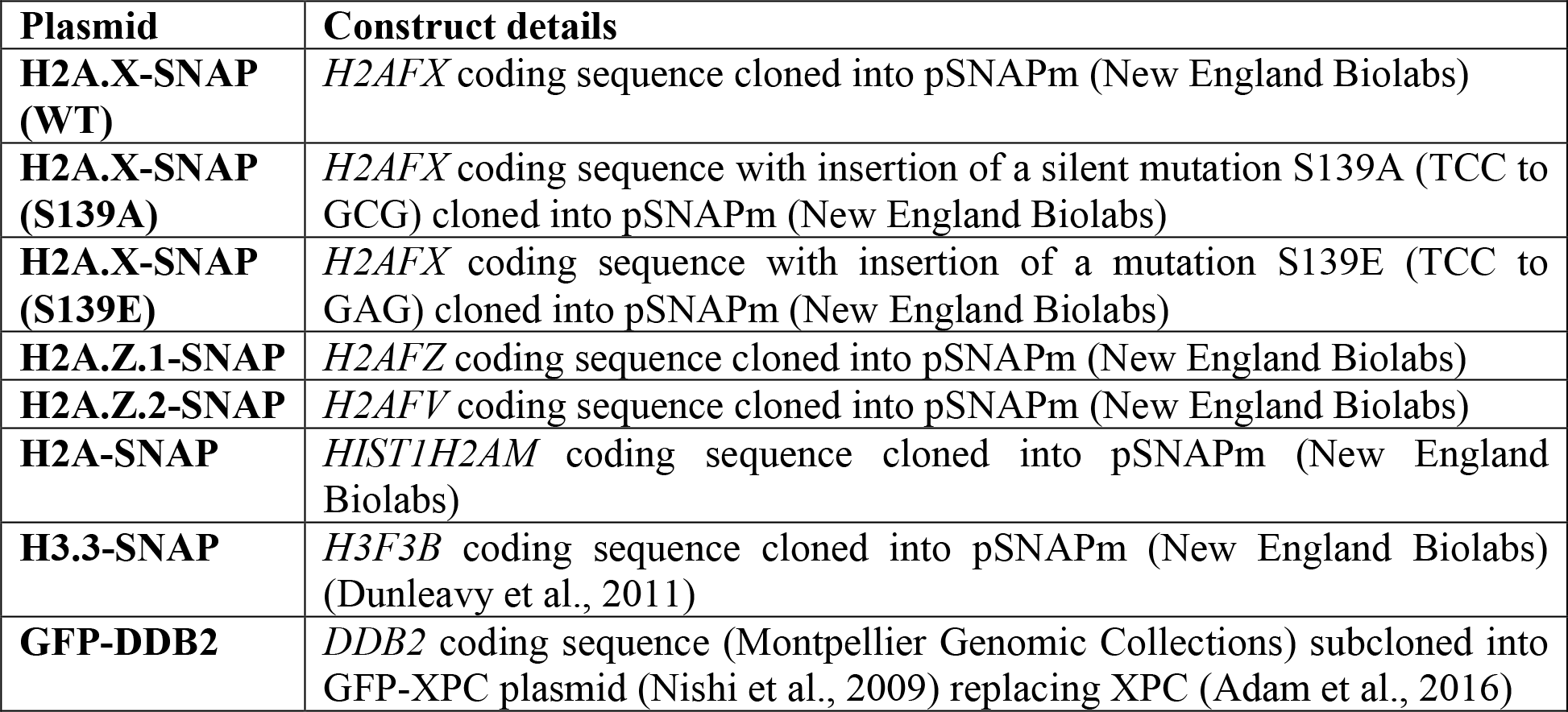
Plasmids. All the coding sequences for histone variants and repair factors are of human origin.

### UVC irradiation

Cells grown on glass coverslips (Menzel Gläser) were irradiated with UVC (254 nm) using a low-pressure mercury lamp. Conditions were set using a VLX-3W dosimeter (Vilbert-Lourmat). For global UVC irradiation, cells in Phosphate Buffer Saline (PBS) were exposed to UVC doses ranging from 4 to 12 J/m^2^ by varying the duration of exposure. For local UVC irradiation (Katsumi et al., 2001; Moné et al., 2001), cells were covered with a polycarbonate filter (5 µm pore size, Millipore) and irradiated with 150 J/m^2^ UVC unless indicated otherwise. Only 2% of the UVC light goes through these micropore filters.

### UVA laser micro-irradiation

Cells grown on glass coverslips (Menzel Gläser) were presensitized with 10 µM 5-bromo-2’-deoxyuridine (BrdU, Sigma-Aldrich) for 24 hr at 37°C. Damage was introduced with a 405 nm laser diode (3 mW) focused through a Plan-Apochromat 63x /1.4 oil objective to yield a spot size of 0.5-1 µm using a LSM710 NLO confocal microscope (Zeiss) and the following laser settings: 40% power, 50 iterations, scan speed 12.6 µsec/pixel.

### Cell extracts and western blot

Total extracts were obtained by scraping cells in Laemmli buffer (50 mM Tris HCl pH 6.8, 1.6% SDS (Sodium Dodecyl Sulfate), 8% glycerol, 4% β-mercaptoethanol, 0.0025% bromophenol blue) followed by 5 min denaturation at 95°C. Cytosolic and nuclear extracts were obtained as previously described (Martini et al., 1998). The chromatin fraction was prepared by addition of benzonase (Novagen) to the pellet after nuclear extraction.

For western blot analysis, extracts were run on 4%–20% Mini-PROTEAN TGX gels (Bio-Rad) in running buffer (200 mM glycine, 25 mM Tris, 0.1% SDS) and transferred onto nitrocellulose membranes (Amersham) with a Trans-Blot SD semidry transfer cell (Bio-Rad). Total proteins were revealed by reversible protein stain (Pierce). Proteins of interest were probed using the appropriate primary and HRP (Horse Radish Peroxidase)-conjugated secondary antibodies (Jackson Immunoresearch), detected using Super-Signal West Pico or Femto chemi-luminescence substrates (Pierce). Alternatively, when fluorescence detection was used instead of chemi-luminescence, total proteins were revealed with REVERT total protein stain, secondary antibodies were conjugated to IRDye 680RD or 800CW and imaging was performed with Odyssey Fc-imager (LI-COR Biosciences) (see Table 4 for the list of antibodies).

**Table 4.**
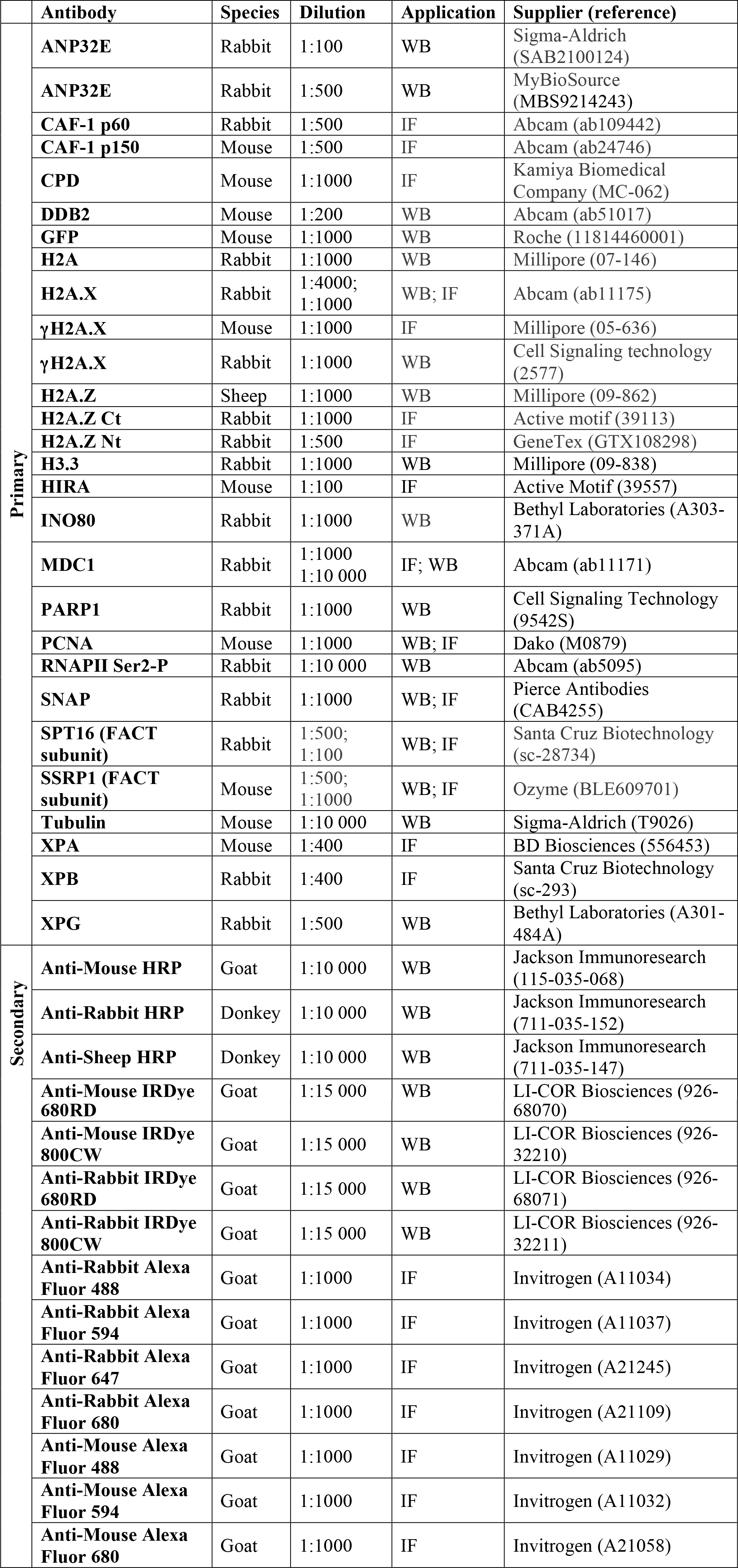
Antibodies.

### Immunofluorescence, image acquisition and analysis

Cells grown on sterile round glass coverslips 12 mm diameter, thickness No.1.5 (Menzel Gläser) were either fixed directly with 2% paraformaldehyde (PFA, Electron Microscopy Sciences) and permeabilized with 0.5% Triton X-100 in PBS or first pre-extracted with 0.5% Triton X-100 in CSK buffer (Cytoskeletal buffer: 10 mM PIPES pH 7.0, 100 mM NaCl, 300 mM sucrose, 3 mM MgCl_2_) and then fixed with 2% PFA to remove soluble proteins (not bound to chromatin). For CPD staining, DNA was denatured with 0.5 M NaOH for 5 min. For PCNA staining, cells were treated with Methanol (Prolabo) for 15 min at −20°C. Samples were blocked in 5% BSA (Bovine Serum Albumin, Sigma-Aldrich) in PBS supplemented with 0.1% Tween 20 (Euromedex) before incubation with primary antibodies and secondary antibodies conjugated to Alexa-Fluor 488, 594, 647 or 680 (Invitrogen) (see Table 4 for a description of the antibodies).

Coverslips were mounted in Vectashield medium with DAPI (Vector Laboratories) and observed with a Leica DMI6000 epifluorescence microscope using a 40x or 63x oil objective. Images were captured using a CCD camera (Hamamatsu) and Metamorph software. For confocal images, we used a Zeiss LSM710 confocal microscope with a Plan-Apochromat 63x/1.4 oil objective and Zen software. Image J (U. S. National Institutes of Health, Bethesda, Maryland, USA, http://imagej.nih.gov/ij/) and ICY (Institut Pasteur, Paris, France, icy.bioimageanalysis.org/) softwares were used for image analysis.

### Visualization of Replicative and Repair Synthesis

To visualize replicative synthesis, EdU (Ethynyl-deoxyUridine) was incorporated into cells during 15 min (10 µM final concentration) and revealed using the Click-iT EdU Alexa Fluor 594 (or 647) Imaging kit (Invitrogen) according to manufacturer’s instructions. To visualize repair synthesis, EdU (10 µM final concentration) was incorporated into cells during 4 h immediately after local UVC irradiation and revealed using the same kit, before CPD labeling by immunofluorescence.

### Nascent RNA labeling

Cells were incubated in DMEM supplemented with 0.5 mM EU (EthynylUridine) for 45 min, rinsed in cold medium and in PBS before fixation in 2% paraformaldehyde. EU incorporation was revealed with Click-iT RNA Imaging kits (Invitrogen) using Alexa Fluor 594 dye according to manufacturer’s instructions.

### Flow cytometry

Cells were fixed in ice-cold 70% ethanol before DNA staining with 50 µg/mL propidium iodide (Sigma-Aldrich) in PBS containing 0.05% Tween 20 and 0.5 mg/mL RNaseA (USB/Affymetrix). DNA content was analyzed by flow cytometry using a FACS Calibur Flow Cytometer (BD Biosciences) and FlowJo Software (TreeStar).

### Colony-Forming Assays

Cells were replated 48 h after siRNA transfection and exposed to global UVC irradiation (4, 8 and 12 J/m^2^) the following day. Colonies were stained 10 days later with 0.5% crystal violet/20% ethanol and counted. Results were normalized to plating efficiencies.

### Statistical analyses

Percentages of positively stained cells were obtained by scoring at least 100 cells in each experiment unless indicated otherwise. In experiments using transcription inhibitors, only cells showing above background TMR levels were scored. Statistical tests were performed using GraphPad Prism. Loss or enrichment of H2A.Z at UV sites relative to the whole nucleus was compared to a theoretical mean of 1 by one-sample t-tests. P-values for mean comparisons between two groups were calculated with a Student’s t-test with Welch’s correction when necessary. Multiple comparisons were performed by one-way ANOVA with Bonferroni post-tests. Comparisons of clonogenic survival and CPD removal kinetics were based on non-linear regression with a polynomial quadratic model. ns: non-significant, *: p < 0.05, **: p < 0.01, ***: p < 0.001.

## AUTHOR CONTRIBUTIONS

S.P., F.L.P., S-K.B. and S.E.P designed and performed experiments and analyzed the data. O.C. provided technical assistance and established mouse cell lines. S.A. generated the H2A.X-SNAP plasmids and the H2A-SNAP and H2A.Z.1-SNAP cell lines. S.E.P. supervised the project and wrote the manuscript with inputs from all authors.

## ACKNOWLEDGMENTS

We thank P-A. Defossez, C. Francastel, V. Mezger and J. Weitzman for critical reading of the manuscript. We thank E. Dunleavy, A. Hamiche and S. Jackson for sharing reagents. We acknowledge the ImagoSeine core facility (Institut Jacques Monod, France BioImaging) for assistance with confocal microscopy. This work was supported by the European Research Council (ERC starting grant ERC-2013-StG-336427 “EpIn”), the French National Research Agency (ANR-12-JSV6-0002-01), the “Who am I?” laboratory of excellence (ANR-11-LABX-0071) funded by the French Government through its “Investments for the Future” program (ANR-11-IDEX-0005-01), EDF Radiobiology program RB 2014-01, the Fondation ARC and France-BioImaging (ANR-10-INSB-04). S.P. is an EMBO Young Investigator. S.A. was recipient of PhD fellowships from University Pierre et Marie Curie and La Ligue contre le Cancer.

## REFERENCES

Abe, T., Sugimura, K., Hosono, Y., Takami, Y., Akita, M., Yoshimura, A., Tada, S., Nakayama, T., Murofushi, H., Okumura, K., Takeda, S., Horikoshi, M., Seki, M., Enomoto, T., 2011. The histone chaperone facilitates chromatin transcription (FACT) protein maintains normal replication fork rates. Journal of Biological Chemistry 286, 30504–30512. doi:10.1074/jbc.M111.264721

Adam, S., Dabin, J., Chevallier, O., Leroy, O., Baldeyron, C., Corpet, A., Lomonte, P., Renaud, O., Almouzni, G., Polo, S.E., 2016. Real-Time Tracking of Parental Histones Reveals Their Contribution to Chromatin Integrity Following DNA Damage. Mol Cell 64, 65–78. doi:10.1016/j.molcel.2016.08.019

Adam, S., Polo, S.E., Almouzni, G., 2013. Transcription recovery after DNA damage requires chromatin priming by the H3.3 histone chaperone HIRA. Cell 155, 94–106. doi:10.1016/j.cell.2013.08.029

Aguilera, A., García-Muse, T., 2013. Causes of genome instability. Annu Rev Genet 47, 1–32. doi:10.1146/annurev-genet-111212-133232

Alabert, C., Barth, T.K., Reverón-Gómez, N., Sidoli, S., Schmidt, A., Jensen, O.N., Imhof, A., Groth, A., 2015. Two distinct modes for propagation of histone PTMs across the cell cycle. Genes Dev 29, 585–590. doi:10.1101/gad.256354.114

Alabert, C., Bukowski-Wills, J.-C., Lee, S.-B., Kustatscher, G., Nakamura, K., de Lima Alves, F., Menard, P., Mejlvang, J., Rappsilber, J., Groth, A., 2014. Nascent chromatin capture proteomics determines chromatin dynamics during DNA replication and identifies unknown fork components. Nat Cell Biol 16, 281–293. doi:10.1038/ncb2918

Alatwi, H.E., Downs, J.A., 2015. Removal of H2A.Z by INO80 promotes homologous recombination. EMBO Rep 16, 986–994. doi:10.15252/embr.201540330

Allis, C.D., Jenuwein, T., 2016. The molecular hallmarks of epigenetic control. Nat Rev Genet 17, 487–500. doi:10.1038/nrg.2016.59

Altmeyer, M., Lukas, J., 2013. To spread or not to spread–chromatin modifications in response to DNA damage. Curr Opin Genet Dev 23, 156–165. doi:10.1016/j.gde.2012.11.001

Atsumi, Y., Minakawa, Y., Ono, M., Dobashi, S., Shinohe, K., Shinohara, A., Takeda, S., Takagi, M., Takamatsu, N., Nakagama, H., Teraoka, H., Yoshioka, K.-I., 2015. ATM and SIRT6/SNF2H Mediate Transient H2AX Stabilization When DSBs Form by Blocking HUWE1 to Allow Efficient γH2AX Foci Formation. Cell Rep 13, 2728–2740. doi:10.1016/j.celrep.2015.11.054

Bannister, A.J., Kouzarides, T., 2011. Regulation of chromatin by histone modifications. Cell Res 21, 381–395. doi:10.1038/cr.2011.22

Bassing, C.H., Chua, K.F., Sekiguchi, J., Suh, H., Whitlow, S.R., Fleming, J.C., Monroe, B.C., Ciccone, D.N., Yan, C., Vlasakova, K., Livingston, D.M., Ferguson, D.O., Scully, R., Alt, F.W., 2002. Increased ionizing radiation sensitivity and genomic instability in the absence of histone H2AX. Proc Natl Acad Sci USA 99, 8173–8178. doi:10.1073/pnas.122228699

Belotserkovskaya, R., Oh, S., Bondarenko, V., Orphanides, G., Studitsky, V., Reinberg, D., 2003. FACT facilitates transcription-dependent nucleosome alteration. Science 301, 1090–1093.

Bensaude, O., 2011. Inhibiting eukaryotic transcription: Which compound to choose? How to evaluate its activity? Transcription 2, 103–108. doi:10.4161/trns.2.3.16172

Bodor, D., Rodríguez, M., Moreno, N., 2012. Analysis of Protein Turnover by Quantitative SNAP‐Based Pulse‐Chase Imaging. Curr Protoc Cell Biol Chapter 8: Unit8.8. doi:10.1002/0471143030.cb0808s55

Bonner, W.M., Redon, C.E., Dickey, J.S., Nakamura, A.J., Sedelnikova, O.A., Solier, S., Pommier, Y., 2008. GammaH2AX and cancer. Nat Rev Cancer 8, 957–967. doi:10.1038/nrc2523

Boyarchuk, E., Filipescu, D., Vassias, I., Cantaloube, S., Almouzni, G., 2014. The histone variant composition of centromeres is controlled by the pericentric heterochromatin state during the cell cycle. J Cell Sci 127, 3347–3359. doi:10.1242/jcs.148189

Buschbeck, M., Hake, S.B., 2017. Variants of core histones and their roles in cell fate decisions, development and cancer. Nat Rev Mol Cell Biol 18, 299–314. doi:10.1038/nrm.2016.166

Chen, C.-C., Carson, J.J., Feser, J., Tamburini, B., Zabaronick, S., Linger, J., Tyler, J.K., 2008. Acetylated lysine 56 on histone H3 drives chromatin assembly after repair and signals for the completion of repair. Cell 134, 231–243. doi:10.1016/j.cell.2008.06.035

Ciccia, A., Elledge, S.J., 2010. The DNA damage response: making it safe to play with knives. Mol Cell 40, 179–204. doi:10.1016/j.molcel.2010.09.019

Dabin, J., Fortuny, A., Polo, S.E., 2016. Epigenome Maintenance in Response to DNA Damage. Mol Cell 62, 712–727. doi:10.1016/j.molcel.2016.04.006

Dantuma, N.P., van Attikum, H., 2016. Spatiotemporal regulation of posttranslational modifications in the DNA damage response. EMBO J 35, 6–23. doi:10.15252/embj.201592595

Diao, L.-T., Chen, C.-C., Dennehey, B., Pal, S., Wang, P., Shen, Z.-J., Deem, A., Tyler, J.K., 2017. Delineation of the role of chromatin assembly and the Rtt101Mms1 E3 ubiquitin ligase in DNA damage checkpoint recovery in budding yeast. PLoS ONE 12, e0180556. doi:10.1371/journal.pone.0180556

Dinant, C., Ampatziadis-Michailidis, G., Lans, H., Tresini, M., Lagarou, A., Grosbart, M., Theil, A.F., van Cappellen, W.A., Kimura, H., Bartek, J., Fousteri, M., Houtsmuller, A.B., Vermeulen, W., Marteijn, J.A., 2013. Enhanced chromatin dynamics by FACT promotes transcriptional restart after UV-induced DNA damage. Mol Cell 51, 469–479. doi:10.1016/j.molcel.2013.08.007

Dunleavy, E.M., Almouzni, G., Karpen, G.H., 2011. H3.3 is deposited at centromeres in S phase as a placeholder for newly assembled CENP-A in G" phase. nucleus 2, 146–157. doi:10.4161/nucl.2.2.15211

Fan, J., Otterlei, M., Wong, H.-K., Tomkinson, A.E., Wilson, D.M., 2004. XRCC1 co-localizes and physically interacts with PCNA. Nucleic Acids Research 32, 2193–2201. doi:10.1093/nar/gkh556

Fan, J.Y., Gordon, F., Luger, K., Hansen, J.C., Tremethick, D.J., 2002. The essential histone variant H2A.Z regulates the equilibrium between different chromatin conformational states. Nat. Struct. Biol. 9, 172–176. doi:10.1038/nsb767

Fan, J.Y., Rangasamy, D., Luger, K., Tremethick, D.J., 2004. H2A.Z alters the nucleosome surface to promote HP1alpha-mediated chromatin fiber folding. Mol Cell 16, 655–661. doi:10.1016/j.molcel.2004.10.023

Gao, Y., Li, C., Wei, L., Teng, Y., Nakajima, S., Chen, X., Xu, J., Legar, B., Ma, H., Spagnol, S.T., Wan, Y., Dahl, K.N., Liu, Y., Levine, A.S., Lan, L., 2017. SSRP1 Cooperates with PARP and XRCC1 to Facilitate Single-Strand DNA Break Repair by Chromatin Priming. Cancer Res 77, 2674–2685. doi:10.1158/0008-5472.CAN-16-3128

Gasparian, A.V., Burkhart, C.A., Purmal, A.A., Brodsky, L., Pal, M., Saranadasa, M., Bosykh, D.A., Commane, M., Guryanova, O.A., Pal, S., Safina, A., Sviridov, S., Koman, I.E., Veith, J., Komar, A.A., Gudkov, A.V., Gurova, K.V., 2011. Curaxins: anticancer compounds that simultaneously suppress NF-κB and activate p53 by targeting FACT. Science translational medicine 3, 95ra74. doi:10.1126/scitranslmed.3002530

Gurard-Levin, Z.A., Quivy, J.-P., Almouzni, G., 2014. Histone chaperones: assisting histone traffic and nucleosome dynamics. Annu Rev Biochem 83, 487–517. doi:10.1146/annurevbiochem-060713-035536

Gursoy-Yuzugullu, O., Ayrapetov, M.K., Price, B.D., 2015. Histone chaperone Anp32e removes H2A.Z from DNA double-strand breaks and promotes nucleosome reorganization and DNA repair. Proc Natl Acad Sci USA 112, 7507–7512. doi:10.1073/pnas.1504868112

Hammond, C.M., Strømme, C.B., Huang, H., Patel, D.J., Groth, A., 2017. Histone chaperone networks shaping chromatin function. Nat Rev Mol Cell Biol 18, 141–158. doi:10.1038/nrm.2016.159

Hanasoge, S., Ljungman, M., 2007. H2AX phosphorylation after UV irradiation is triggered by DNA repair intermediates and is mediated by the ATR kinase. Carcinogenesis 28, 2298–2304. doi:10.1093/carcin/bgm157

Heo, K., Kim, H., Choi, S.H., Choi, J., Kim, K., Gu, J., Lieber, M.R., Yang, A.S., An, W., 2008. FACT-mediated exchange of histone variant H2AX regulated by phosphorylation of H2AX and ADP-ribosylation of Spt16. Mol Cell 30, 86–97. doi:10.1016/j.molcel.2008.02.029

Herrera-Moyano, E., Mergui, X., García-Rubio, M.L., Barroso, S., Aguilera, A., 2014. The yeast and human FACT chromatin-reorganizing complexes solve R-loop-mediated transcription-replication conflicts. Genes Dev 28, 735–748. doi:10.1101/gad.234070.113

Hoeijmakers, J.H.J., 2009. DNA damage, aging, and cancer. The New England journal of medicine 361, 1475–1485. doi:10.1056/NEJMra0804615

Jackson, S.P., Bartek, J., 2009. The DNA-damage response in human biology and disease. Nature 461, 1071–1078. doi:10.1038/nature08467

Jackson, V., 1990. In vivo studies on the dynamics of histone-DNA interaction: evidence for nucleosome dissolution during replication and transcription and a low level of dissolution independent of both. Biochemistry 29, 719–731.

Jackson, V., 1987. Deposition of newly synthesized histones: new histones H2A and H2B do not deposit in the same nucleosome with new histones H3 and H4. Biochemistry 26, 2315–2325.

Jeronimo, C., Watanabe, S., Kaplan, C.D., Peterson, C.L., Robert, F., 2015. The Histone Chaperones FACT and Spt6 Restrict H2A.Z from Intragenic Locations. Mol Cell 58, 1113–1123. doi:10.1016/j.molcel.2015.03.030

Kim, J., Sturgill, D., Sebastian, R., Khurana, S., Tran, A.D., Edwards, G.B., Kruswick, A., Burkett, S., Hosogane, E.K., Hannon, W.W., Weyemi, U., Bonner, W.M., Luger, K., Oberdoerffer, P., 2018. Replication Stress Shapes a Protective Chromatin Environment across Fragile Genomic Regions. Mol Cell 69, 36–47. doi:10.1016/j.molcel.2017.11.021

Kim, J.-A., Haber, J.E., 2009. Chromatin assembly factors Asf1 and CAF-1 have overlapping roles in deactivating the DNA damage checkpoint when DNA repair is complete. Proc Natl Acad Sci U S A 106, 1151–1156. doi:10.1073/pnas.0812578106

Kimura, H., Cook, P.R., 2001. Kinetics of core histones in living human cells: little exchange of H3 and H4 and some rapid exchange of H2B. J Cell Biol 153, 1341–1353.

Kurat, C.F., Yeeles, J.T.P., Patel, H., Early, A., Diffley, J.F.X., 2017. Chromatin Controls DNA Replication Origin Selection, Lagging-Strand Synthesis, and Replication Fork Rates. Mol Cell 65, 117–130. doi:10.1016/j.molcel.2016.11.016

Lazzaro, F., Giannattasio, M., Puddu, F., Granata, M., Pellicioli, A., Plevani, P., Muzi Falconi, M., 2009. Checkpoint mechanisms at the intersection between DNA damage and repair. DNA Rep 8, 1055–1067. doi:10.1016/j.dnarep.2009.04.022

Liu, Y., Parry, J.A., Chin, A., Duensing, S., Duensing, A., 2008. Soluble histone H2AX is induced by DNA replication stress and sensitizes cells to undergo apoptosis. Molecular cancer 7, 61. doi:10.1186/1476-4598-7-61

Louters, L., Chalkley, R., 1985. Exchange of histones H1, H2A, and H2B in vivo. Biochemistry 24, 3080–3085.

Luger, K., Dechassa, M.L., Tremethick, D.J., 2012. New insights into nucleosome and chromatin structure: an ordered state or a disordered affair? Nat Rev Mol Cell Biol 13, 436–447. doi:10.1038/nrm3382

Luijsterburg, M.S., de Krijger, I., Wiegant, W.W., Shah, R.G., Smeenk, G., de Groot, A.J.L., Pines, A., Vertegaal, A.C.O., Jacobs, J.J.L., Shah, G.M., van Attikum, H., 2016. PARP1 Links CHD2-Mediated Chromatin Expansion and H3.3 Deposition to DNA Repair by Non-homologous End-Joining. Mol Cell 61, 547–562. doi:10.1016/j.molcel.2016.01.019

Luijsterburg, M.S., Lindh, M., Acs, K., Vrouwe, M.G., Pines, A., van Attikum, H., Mullenders, L.H., Dantuma, N.P., 2012. DDB2 promotes chromatin decondensation at UV-induced DNA damage. J Cell Biol 197, 267–281. doi:10.1083/jcb.201106074

Mailand, N., Gibbs-Seymour, I., Bekker-Jensen, S., 2013. Regulation of PCNA-protein interactions for genome stability. Nat Rev Mol Cell Biol 14, 269–282. doi:10.1038/nrm3562

Mannironi, C., Bonner, W.M., Hatch, C.L., 1989. H2A.X. a histone isoprotein with a conserved C-terminal sequence, is encoded by a novel mRNA with both DNA replication type and polyA 3′ processing signals. Nucleic Acids Research 17, 9113–9126.

Mao, Z., Pan, L., Wang, W., Sun, J., Shan, S., Dong, Q., Liang, X., Dai, L., Ding, X., Chen, S., Zhang, Z., Zhu, B., Zhou, Z., 2014. Anp32e, a higher eukaryotic histone chaperone directs preferential recognition for H2A.Z. Cell Res 24, 389–399. doi:10.1038/cr.2014.30

Marini, F., Nardo, T., Giannattasio, M., Minuzzo, M., Stefanini, M., Plevani, P., Muzi Falconi, M., 2006. DNA nucleotide excision repair-dependent signaling to checkpoint activation. Proc Natl Acad Sci USA 103, 17325–17330. doi:10.1073/pnas.0605446103

Marteijn, J.A., Bekker-Jensen, S., Mailand, N., Lans, H., Schwertman, P., Gourdin, A.M., Dantuma, N.P., Lukas, J., Vermeulen, W., 2009. Nucleotide excision repair-induced H2A ubiquitination is dependent on MDC1 and RNF8 and reveals a universal DNA damage response. J Cell Biol 186, 835–847. doi:10.1083/jcb.200902150

Moggs, J.G., Grandi, P., Quivy, J.P., Jónsson, Z.O., Hübscher, U., Becker, P.B., Almouzni, G., 2000. A CAF-1-PCNA-mediated chromatin assembly pathway triggered by sensing DNA damage. Molecular and Cellular Biology 20, 1206–1218.

Moser, J., Kool, H., Giakzidis, I., Caldecott, K., Mullenders, L.H.F., Fousteri, M.I., 2007. Sealing of chromosomal DNA nicks during nucleotide excision repair requires XRCC1 and DNA ligase III alpha in a cell-cycle-specific manner. Mol Cell 27, 311–323. doi:10.1016/j.molcel.2007.06.014

Nishi, R., Alekseev, S., Dinant, C., Hoogstraten, D., Houtsmuller, A.B., Hoeijmakers, J.H.J., Vermeulen, W., Hanaoka, F., Sugasawa, K., 2009. UV-DDB-dependent regulation of nucleotide excision repair kinetics in living cells. DNA Repair (Amst) 8, 767–776. doi:10.1016/j.dnarep.2009.02.004

Nishibuchi, I., Suzuki, H., Kinomura, A., Sun, J., Liu, N.-A., Horikoshi, Y., Shima, H., Kusakabe, M., Harata, M., Fukagawa, T., Ikura, T., Ishida, T., Nagata, Y., Tashiro, S., 2014. Reorganization of damaged chromatin by the exchange of histone variant H2A.Z-2. Int. J. Radiat. Oncol. Biol. Phys. 89, 736–744. doi:10.1016/j.ijrobp.2014.03.031

Obri, A., Ouararhni, K., Papin, C., Diebold, M.-L., Padmanabhan, K., Marek, M., Stoll, I., Roy, L., Reilly, P.T., Mak, T.W., Dimitrov, S., Romier, C., Hamiche, A., 2014. ANP32E is a histone chaperone that removes H2A.Z from chromatin. Nature 505, 648–653. doi:10.1038/nature12922

Okuhara, K., Ohta, K., Seo, H., Shioda, M., Yamada, T., Tanaka, Y., Dohmae, N., Seyama, Y., Shibata, T., Murofushi, H., 1999. A DNA unwinding factor involved in DNA replication in cell-free extracts of Xenopus eggs. Curr Biol 9, 341–350.

Papamichos-Chronakis, M., Krebs, J.E., Peterson, C.L., 2006. Interplay between Ino80 and Swr1 chromatin remodeling enzymes regulates cell cycle checkpoint adaptation in response to DNA damage. Genes Dev 20, 2437–2449. doi:10.1101/gad.1440206

Polo, S.E., Almouzni, G., 2015. Chromatin dynamics after DNA damage: The legacy of the access-repair-restore model. DNA Repair (Amst) 36, 114–121. doi:10.1016/j.dnarep.2015.09.014

Polo, S.E., Roche, D., Almouzni, G., 2006. New histone incorporation marks sites of UV repair in human cells. Cell 127, 481–493. doi:10.1016/j.cell.2006.08.049

Ray-Gallet, D., Woolfe, A., Vassias, I., Pellentz, C., Lacoste, N., Puri, A., Schultz, D.C., Pchelintsev, N.A., Adams, P.D., Jansen, L.E.T., Almouzni, G., 2011. Dynamics of histone h3 deposition in vivo reveal a nucleosome gap-filling mechanism for h3.3 to maintain chromatin integrity. Mol Cell 44, 928–941. doi:10.1016/j.molcel.2011.12.006

Rocha, C.R.R., Lerner, L.K., Okamoto, O.K., Marchetto, M.C., Menck, C.F.M., 2013. The role of DNA repair in the pluripotency and differentiation of human stem cells. Mutat Res 752, 25–35. doi:10.1016/j.mrrev.2012.09.001

Rogakou, E.P., Boon, C., Redon, C., Bonner, W.M., 1999. Megabase chromatin domains involved in DNA double-strand breaks in vivo. J Cell Biol 146, 905–916.

Rogakou, E.P., Pilch, D.R., Orr, A.H., Ivanova, V.S., Bonner, W.M., 1998. DNA double-stranded breaks induce histone H2AX phosphorylation on serine 139. J Biol Chem 273, 5858–5868.

Safina, A., Cheney, P., Pal, M., Brodsky, L., Ivanov, A., Kirsanov, K., Lesovaya, E., Naberezhnov, D., Nesher, E., Koman, I., Wang, D., Wang, J., Yakubovskaya, M., Winkler, D., Gurova, K., 2017. FACT is a sensor of DNA torsional stress in eukaryotic cells. Nucleic Acids Research 45, 1925–1945. doi:10.1093/nar/gkw1366

Seo, J., Kim, K., Chang, D.-Y., Kang, H.-B., Shin, E.-C., Kwon, J., Choi, J.K., 2014. Genome-wide reorganization of histone H2AX toward particular fragile sites on cell activation. Nucleic Acids Research 42, 1016–1025. doi:10.1093/nar/gkt951

Seo, J., Kim, S.C., Lee, H.-S., Kim, J.K., Shon, H.J., Salleh, N.L.M., Desai, K.V., Lee, J.H., Kang, E.-S., Kim, J.S., Choi, J.K., 2012. Genome-wide profiles of H2AX and γ-H2AX differentiate endogenous and exogenous DNA damage hotspots in human cells. Nucleic Acids Research 40, 5965–5974. doi:10.1093/nar/gks287

Shaltiel, I.A., Krenning, L., Bruinsma, W., Medema, R.H., 2015. The same, only different - DNA damage checkpoints and their reversal throughout the cell cycle. J Cell Sci 128, 607–620. doi:10.1242/jcs.163766

Shen, X., Mizuguchi, G., Hamiche, A., Wu, C., 2000. A chromatin remodelling complex involved in transcription and DNA processing. Nature 406, 541–544. doi:10.1038/35020123

Smeenk, G., van Attikum, H., 2013. The chromatin response to DNA breaks: leaving a mark on genome integrity. Annu Rev Biochem 82, 55–80. doi:10.1146/annurev-biochem-061809-174504

Smerdon, M.J., 1991. DNA repair and the role of chromatin structure. Curr Opin Cell Biol 3, 422–428.

Talbert, P.B., Henikoff, S., 2017. Histone variants on the move: substrates for chromatin dynamics. Nat Rev Mol Cell Biol 18, 115–126. doi:10.1038/nrm.2016.148

Talbert, P.B., Henikoff, S., 2010. Histone variants–ancient wrap artists of the epigenome. Nat Rev Mol Cell Biol 11, 264–275. doi:10.1038/nrm2861

Tan, B., Chien, C., Hirose, S., Lee, S., 2006. Functional cooperation between FACT and MCM helicase facilitates initiation of chromatin DNA replication. EMBO J 25, 3975–3985.

Weyemi, U., Redon, C.E., Choudhuri, R., Aziz, T., Maeda, D., Boufraqech, M., Parekh, P.R., Sethi, T.K., Kasoji, M., Abrams, N., Merchant, A., Rajapakse, V.N., Bonner, W.M., 2016. The histone variant H2A.X is a regulator of the epithelial-mesenchymal transition. Nat Commun 7, 10711. doi:10.1038/ncomms10711

Wu, R.S., Bonner, W.M., 1981. Separation of basal histone synthesis from S-phase histone synthesis in dividing cells. Cell 27, 321–330.

Wu, T., Liu, Y., Wen, D., Tseng, Z., Tahmasian, M., Zhong, M., Rafii, S., Stadtfeld, M., Hochedlinger, K., Xiao, A., 2014. Histone variant H2A.X deposition pattern serves as a functional epigenetic mark for distinguishing the developmental potentials of iPSCs. Cell Stem Cell 15, 281–294. doi:10.1016/j.stem.2014.06.004

Xu, Y., Ayrapetov, M.K., Xu, C., Gursoy-Yuzugullu, O., Hu, Y., Price, B.D., 2012. Histone H2A.Z controls a critical chromatin remodeling step required for DNA double-strand break repair. Mol Cell 48, 723–733. doi:10.1016/j.molcel.2012.09.026

Yang, J., Zhang, X., Feng, J., Leng, H., Li, S., Xiao, J., Liu, S., Xu, Z., Xu, J., Li, D., Wang, Z., Wang, J., Li, Q., 2016. The Histone Chaperone FACT Contributes to DNA Replication-Coupled Nucleosome Assembly. Cell Rep 14, 1128–1141. doi:10.1016/j.celrep.2015.12.096

Zhang, H., Roberts, D.N., Cairns, B.R., 2005. Genome-wide dynamics of Htz1, a histone H2A variant that poises repressed/basal promoters for activation through histone loss. Cell 123, 219–231. doi:10.1016/j.cell.2005.08.036

Zhou, C.Y., Johnson, S.L., Gamarra, N.I., Narlikar, G.J., 2016. Mechanisms of ATP-Dependent Chromatin Remodeling Motors. Annu Rev Biophys 45, 153–181. doi:10.1146/annurev-biophys-051013-022819

